# The carnitine shuttle links mitochondrial metabolism to histone acetylation and lipogenesis

**DOI:** 10.1101/2022.09.24.509197

**Authors:** Luke Izzo, Sophie Trefely, Christina Demetriadou, Jack Drummond, Takuya Mizukami, Nina Kuprasertkul, Aimee Farria, Phuong Nguyen, Lauren Reich, Joshua Shaffer, Hayley Affronti, Alessandro Carrer, Andrew Andrews, Brian C. Capell, Nathaniel W. Snyder, Kathryn E. Wellen

## Abstract

Acetyl-CoA is a central metabolite used for lipid synthesis in the cytosol and histone acetylation in the nucleus, among other pathways. The two major precursors to acetyl-CoA in the nuclear-cytoplasmic compartment are citrate and acetate, which are processed to acetyl-CoA by ATP-citrate lyase (ACLY) and acyl-CoA synthetase short-chain 2 (ACSS2), respectively. While some evidence has suggested the existence of additional routes to nuclear-cytosolic acetyl-CoA, such pathways remain poorly defined. To investigate this, we generated cancer cell lines lacking both ACLY and ACSS2. Unexpectedly, and in contrast to observations in fibroblasts, ACLY and ACSS2 double knockout (DKO) cancer cells remain viable and proliferate, maintain pools of cytosolic acetyl-CoA, and are competent to acetylate proteins in both cytosolic and nuclear compartments. Using stable isotope tracing, we show that both glucose and fatty acids feed acetyl-CoA pools and histone acetylation in DKO cells. Moreover, we provide evidence for the carnitine shuttle and carnitine acetyltransferase (CrAT) as a substantial pathway to transfer two-carbon units from mitochondria to cytosol independent of ACLY. Indeed, in the absence of ACLY, glucose can feed fatty acid synthesis in a carnitine responsive and CrAT-dependent manner. This work defines a carnitine-facilitated route to produce nuclear-cytosolic acetyl-CoA, shedding light on the intricate regulation and compartmentalization of acetyl-CoA metabolism

## INTRODUCTION

Acetyl-CoA is a central metabolic intermediate that is generated during nutrient catabolism, used as a building block for lipid synthesis, and serves as the substrate for protein and metabolite acetylation. In mitochondria, acetyl-CoA is generated from the breakdown of nutrients including carbohydrates, fatty acids, and amino acids. Mitochondrial acetyl-CoA enters the tricarboxylic acid cycle through a condensation reaction with oxaloacetate to generate citrate. The mitochondrial pool of acetyl-CoA is spatially distinct from acetyl-CoA found in the nucleus and cytosol due to the inability of acetyl-CoA to cross the inner mitochondrial membrane. Due to this compartmentalization, acetyl-CoA must be generated separately within the nucleus or cytosol for its use in these compartments. This is accomplished by ATP-citrate lyase (ACLY), which cleaves citrate exported from mitochondria into acetyl-CoA and oxaloacetate, and acyl-CoA short chain fatty acid synthase 2 (ACSS2), which condenses acetate with free coenzyme A. Acetyl-CoA generated by these enzymes is used for *de novo* lipogenesis, as well as for acetylation in the nucleus and cytosol^1–3^.

We previously demonstrated that in the absence of ACLY, ACSS2 is upregulated and acetate feeds acetyl-CoA pools for lipogenesis and histone acetylation^4^. Furthermore, *Acly*^-/-^ mouse embryonic fibroblasts (MEFs) are dependent on exogenous acetate for viability, suggesting that acetyl-CoA synthesis by ACSS2 is the primary compensatory mechanism in the absence of ACLY^4^. Such compensation from acetate via ACSS2 is also observed *in vivo* upon deletion of *Acly* from adipose tissue or liver^4–6^. Yet, several clues suggested that additional acetyl-CoA generating mechanisms within the nuclear-cytosolic compartment must exist. First, in the absence of ACLY, about 20-40% of the acetyl-CoA pool in whole cells and in the cytosol remains unlabeled from exogenous acetate^4,7^. Secondly, although the downstream mevalonate pathway intermediate HMG-CoA is rapidly depleted in the absence of ACLY and acetate, acetyl-CoA levels decrease more moderately upon acute acetate withdrawal^8^. Third, histone acetylation is suppressed in the absence of ACLY at physiological acetate concentrations but does not appear to decline further when acetate is withdrawn^4^. Finally, though glucose use for fatty acid synthesis is strongly suppressed in the absence of ACLY, it is not fully blocked *in vitro* or *in vivo*^4,6^. Based on these clues, we hypothesized that two broad mechanisms could potentially account for these findings: 1) intracellular acetate production from other nutrients; and/or 2) another acetyl-CoA producer other than ACLY and ACSS2 (Figure 1A).

**Figure 1:**
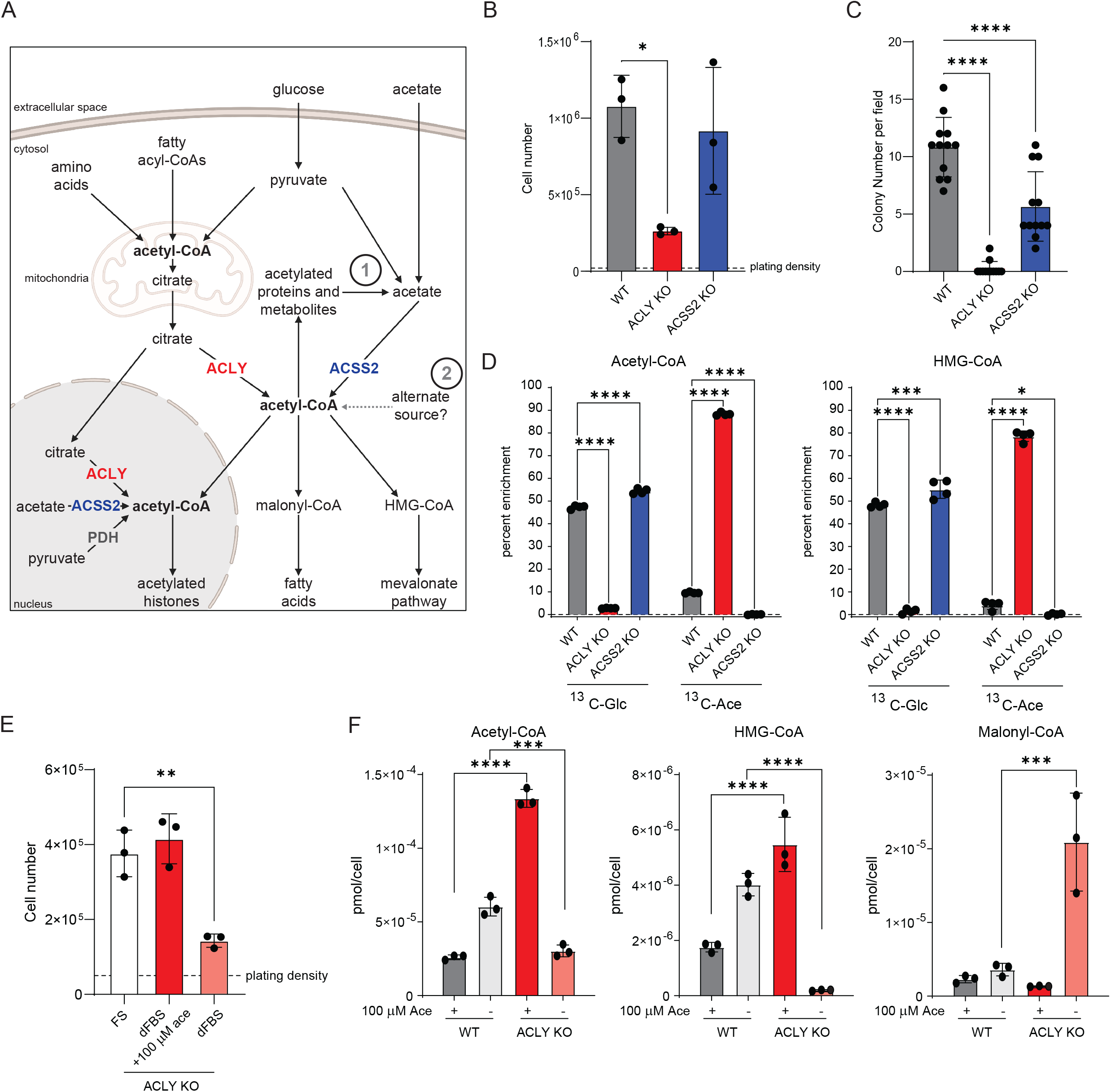
ACLY KO cancer cells proliferate in the absence of exogenous acetate. A) Schematic diagram of acetyl-CoA metabolism. Arrows represent biochemical conversions. Numbers refer to potential ACLY- and exogenous acetate-independent acetyl-CoA generating pathways in the nuclear-cytosolic compartment. Created with BioRender.com. B) Proliferation of WT, ACLY KO, and ACSS2 KO HCC cell lines over 4 days in DMEM + 10% FS. Statistical significance was calculated by one-way ANOVA. C) Colony formation in soft agar of WT, ACLY KO, and ACSS2 KO HCC cells. Colonies were counted at 4× magnification. Data are from 3 replicate wells with 4 fields counted per well. Statistical significance was calculated by one-way ANOVA. D) Whole cell acetyl-CoA measurements and isotopologue distribution (labeled acetyl-CoA, m+2; labeled HMG-CoA sum of m+2, m+4, and m+6) of WT, ACLY KO, and ACSS2 KO HCC cells cultured in glucose and glutamine free DMEM + 10% dFBS + 10mM glucose + 100μM acetate for 6 hours. Statistical significance was calculated by two-way ANOVA. E) ACLY KO proliferation in DMEM + 10% FS, DMEM + 10% dFBS, or DMEM + 10% dFBS + 100μM acetate for 96 hours. Statistical significance was calculated by one-way ANOVA. F) Whole cell acyl-CoA measurements of WT and ACLY KO cells grown in DMEM + 10% dFBS or DMEM + 10% dFBS + 100μM acetate for 24 hours. Statistical significance was calculated by two-way ANOVA. FS, full serum (10% calf serum); dFBS, dialyzed FBS. Each point represents a biological replicate and error bars represent standard deviation. *p≤0.05; **p≤0.01; ***p≤0.001; ****p≤0.0001.

Both types of mechanisms have been reported, but the significance of such pathways remains poorly understood. In terms of endogenous sources, acetate production can occur directly from pyruvate non-enzymatically or via the pyruvate dehydrogenase complex (PDC)^9,10^. Additionally, acetate can be released from histone deacetylation^11^ and acetylated metabolite hydrolysis^12–14^. Furthermore, non-canonical routes to acetyl-CoA production outside of mitochondria have also been proposed, with several studies reporting a moonlighting function of the pyruvate dehydrogenase complex (PDC), which can translocate to the nucleus under specific conditions to generate a local source of acetyl-CoA from pyruvate for histone acetylation^15–18^. Beyond pyruvate to acetate conversion and nuclear PDC, less well understood routes of nuclear-cytosolic acetyl-CoA metabolism have been suggested, including peroxisomal production and export of acetyl-CoA and acetylcarnitine shuttling out of the mitochondria^19–21^. The functional significance of such pathways relative to the canonical pathways via ACLY and ACSS2 remains poorly understood.

To evaluate potential alternative acetyl-CoA producing pathways, we generated cancer cell lines in which ACLY and ACSS2 are genetically deleted individually or in combination. Intriguingly, we show that cancer cells lacking both ACLY and ACSS2 (DKO cells) are viable, proliferate, contain a nuclear-cytosolic pool of acetyl-CoA, and sustain protein acetylation. Using carbon-13 tracing experiments, we demonstrate that fatty acids and glucose are prominent sources of acetyl-CoA that can feed acetyl-CoA pools and histone acetylation in the absence of ACLY and ACSS2. The data indicate that this is mediated at least in part via the carnitine shuttle and carnitine acetyltransferase (CrAT). Carnitine and CrAT function to transport acetyl-units out of the mitochondria and enable glucose dependent acetyl-CoA synthesis and *de novo* lipogenesis in an ACLY-independent manner. Overall, the data demonstrate that ACLY and ACSS2 are not the sole sources of acetyl-CoA in the nuclear-cytosolic compartment, and that the carnitine shuttle participates in the transit of acetyl-CoA from mitochondria to cytosol to support lipogenesis and histone acetylation.

## RESULTS

### Cancer cells maintain viability and proliferation in the absence of ACLY and exogenous acetate

Since cancer cells are adept at engaging available metabolic flexibility mechanisms, we hypothesized that non-canonical acetyl-CoA production mechanisms could be revealed by developing ACLY KO cancer cell models that are not dependent on acetate for viability. We used murine *Acly^flox/flox^* hepatocellular carcinoma (HCC) cell lines and generated isogenic cell lines lacking ACLY or ACSS2 via adenoviral Cre treatment or CRISPR/Cas9 gene editing, respectively (Supplemental Figure 1A). ACLY KO cells had a dramatically decreased proliferation rate in cell growth and soft agar colony formation assays (Figure 1B, C). ACSS2 KO cells had no impairment of 2D cell growth but showed a modest defect in the numbers and size of soft agar colonies formed compared to parental controls (Figure 1B, C; Supplemental Figure 1B). As expected, loss of ACLY potently suppressed uniformly labeled ^13^C_6_-glucose (^13^C-Glc) and increased ^13^C_2_-acetate (^13^C-Ace) incorporation into acetyl-CoA and the downstream metabolite HMG-CoA (Figure 1D). ACSS2 KO cells used glucose similarly to their WT counterparts and labeling from acetate was completely ablated, showing that ACSS2 is the primary acyl-CoA synthetase that mediates synthesis of acetyl-CoA from acetate in these cells (Figure 1D). These data functionally validate our KO cell models.

We next tested if ACLY loss rendered these cancer cells reliant on acetate for proliferation and acetyl-CoA synthesis. While proliferation slows in ACLY KO HCC cells in the absence of acetate, the cells retain viability and continue to proliferate in the absence of exogenous acetate, in contrast to ACLY KO MEFs (Figure 1E; Supplemental Figure 1C)^4^. Similar observations were made in another cancer cell line, derived from Kras^G12D^-driven pancreatic cancer in mice with *Acly* deletion in the pancreas^22^, which also could proliferate in the absence of ACLY and exogenous acetate (Supplemental Figure 1E). Interestingly, these ACLY KO pancreatic cancer cells showed little to no reliance on exogenous acetate for proliferation, which is similar to that previously observed in ACLY null glioblastoma cells^4^. In the HCC ACLY KO cells, we quantified acetyl-CoA in the presence or absence of acetate, finding that acetyl-CoA abundance is elevated in ACLY KO cells in the presence of acetate and decreases back to that in WT cells in its absence (Figure 1F). In contrast, HMG-CoA is almost entirely depleted upon acetate withdrawal, consistent with that reported previously in ACLY KO cells^8^. Unexpectedly, malonyl-CoA accumulated under these conditions (Figure 1F). Together, these findings indicate that while exogenous acetate feeds acetyl-CoA pools in the absence of ACLY, additional mechanisms must also be available to cells to support proliferation.

### Cancer cells proliferate and maintain a pool of cytosolic acetyl-CoA in the absence of both ACLY and ACSS2

One possible explanation for these data is an endogenous source of acetate. To test this, we asked if ACSS2 is required for viability in ACLY KO cells, using CRISPR/Cas9 gene editing to delete *Acss2* in the ACLY KO HCC and pancreatic ACLY KO cells. Single cell clonal selection revealed that cells lacking both ACLY and ACSS2 (DKO) are viable and proliferate, albeit slower (Figure 2A, B; Supplemental Figure 2A, B, C).

**Figure 2:**
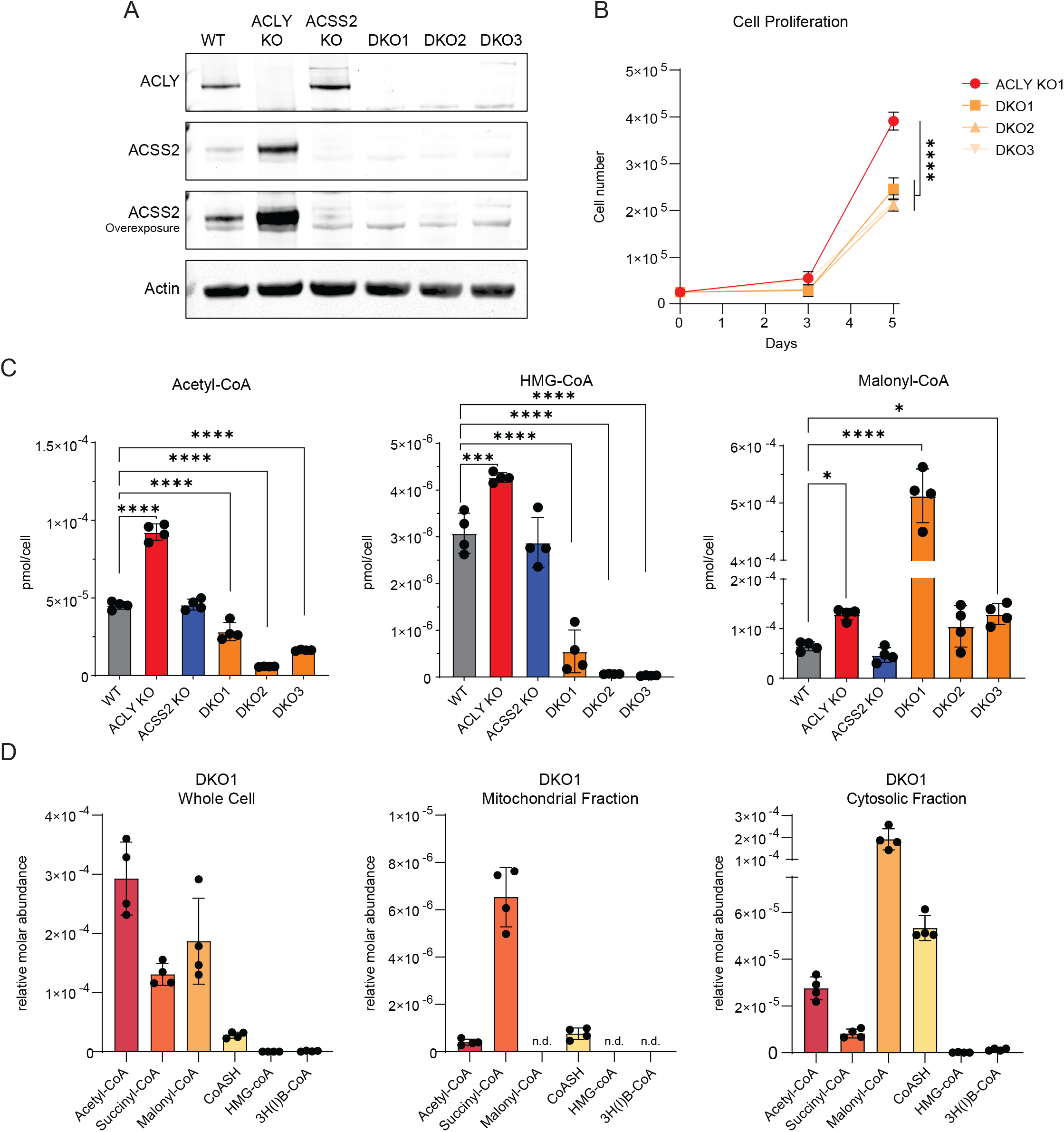
ACLY/ACSS2 double knockout cells are viable and maintain a cytosolic pool of acetyl-CoA. A) Western blot for ACLY and ACSS2 in WT, ACLY KO, ACSS2 KO, and 3 DKO cell lines. B) Proliferation curve of ACLY KO and DKO HCC cell lines over 5 days in DMEM + 10% FS. Data are represented as mean of three replicates +/- standard deviation. Statistical significance was calculated by one-way ANOVA between samples at 5 days. C) Whole cell acyl-CoA measurements in WT, ACLY KO, ACSS2 KO, and DKO cell lines grown in DMEM + 10% dFBS + 100μM acetate for 24 hours. Statistical significance was calculated by one-way ANOVA. D) Acyl-CoA quantitation using SILEC-subcellular fractionation performed on DKO1 cells grown in DMEM + 10% dFBS + 100μM acetate for 24 hours. 3H(I)B-CoA, 3-HB-CoA and isobutyryl-CoA are not resolved and are represented together. Acyl-CoA species marked with n.d. were not detected in n=4 samples. Each point represents a biological replicate and error bars represent standard deviation. *p≤0.05; **p≤0.01; ***p≤0.001; ****p≤0.0001

We next examined acyl-CoA abundance, finding that acetyl-CoA is modestly reduced, HMG-CoA is dramatically reduced, and malonyl-CoA tends to be elevated in DKO cells, similar to the phenotype seen in ACLY KO cells in the absence of acetate (Figure 2C; Supplemental Figure 2D). To determine whether a substantial acetyl-CoA pool is present in the cytosol of DKO cells, we applied SILEC-SF, a recently developed technique for rigorous quantification of acyl-CoAs in subcellular compartments^8^. Subcellular measurements demonstrated distinct acyl-CoA profiles in mitochondria versus cytosol, including a clear cytosolic acetyl-CoA pool in the DKO cells (Figure 2D). The data indicate that cells are able to maintain nuclear-cytosolic acetyl-CoA pools independent of ACLY and ACSS2.

### Cells lacking ACLY and ACSS2 are dependent on exogenous fatty acids

Having established the existence of an extramitochondrial acetyl-CoA pool in the DKO cells, we carried out RNA-sequencing to identify potential compensatory pathways. PCA analysis revealed distinct separation between ACLY deficient (ACLY KO and DKO1) and ACLY proficient genotypes (WT and ACSS2 KO) (Supplemental Figure 3A). Both ACLY KO and DKO cells showed marked transcriptional changes compared to WT with substantial overlap. However, there was also a distinct subset of genes specifically regulated in the DKO cells. (Figure 3A; Supplemental Figure 3B, C; Supplemental Table 1). Thus, we performed gene set enrichment analysis to identify functional groups of genes up- or down-regulated in the DKO cells (Figure 3B). Cell cycle related genes were among those suppressed, while fatty acid metabolism was notably enriched in the DKO cells, including fatty acid oxidation-related genes (Figure 3B, C; Supplemental Figure 3D). These included both mitochondrial and peroxisomal fatty acid oxidation genes (Figure 3C). These data suggest that lipid metabolism may be perturbed in the DKO cells and that fatty acid oxidation pathways might be upregulated as part of a compensatory mechanism.

**Figure 3:**
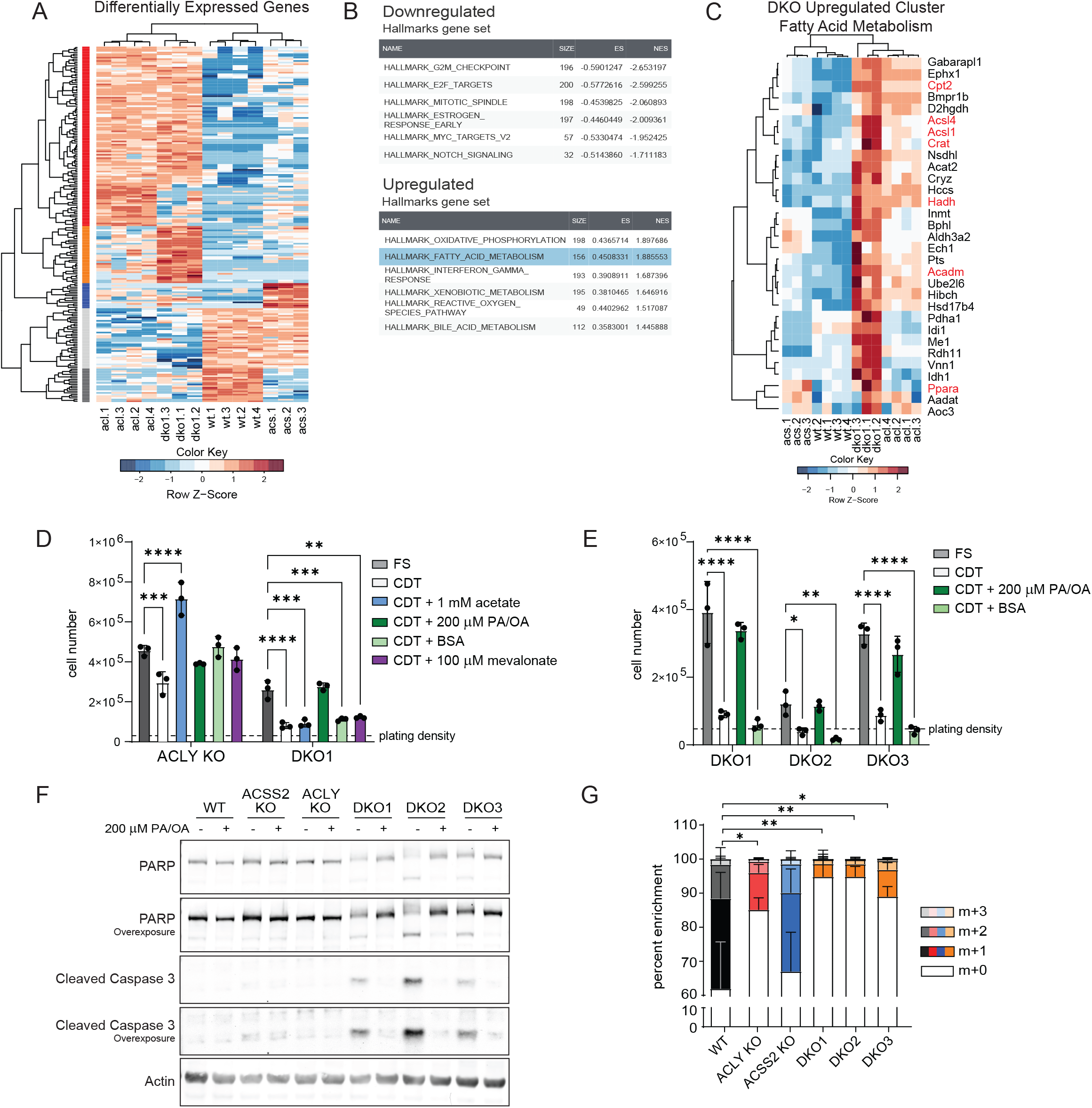
Loss of ACLY and ACSS2 alters fatty acid metabolism and causes reliance on exogenous fatty acids. A) Heatmap of all differentially expressed genes between all 4 genotypes, log2FC>2 and an adjusted p-value<0.01 expressed as row Z-score. DESeq counts were log2 transformed before clustering. Row clusters (red, orange, blue, light gray, and dark gray) represent groups of genes commonly differentially regulated by sample cluster. See Table S1 for gene list by cluster. B) The top six upregulated and downregulated Hallmarks gene sets from GSEA analysis comparing DKO cells to all other genotypes. C) Genes from the hallmarks fatty acid metabolism gene set commonly upregulated across DKO1 samples. Cluster expanded from supplemental figure 3D. DESeq counts were log2 transformed before clustering. D) Cell proliferation after 96 hours. Cells were plated in DMEM/F12 media overnight then cultured in DMEM + 10% FS or CDT serum with or without the addition of metabolites. PA/OA is 100 μM of each fatty acid conjugated to BSA (200 μM total). Statistical significance was calculated by two-way ANOVA. E) Cell proliferation after 96 hours. Cells were plated in DMEM/F12 media overnight then cultured in DMEM + 10% FS or CDT serum with or without the addition of metabolites. PA/OA is 100 μM of each fatty acid conjugated to BSA (200 μM total). BSA condition is equal volume of fatty acid free BSA as added to PA/OA condition. Statistical significance was calculated by two-way ANOVA. F) Western blot analysis of cells cultured in DMEM + 10% CDT serum with or without PA/OA. PA/OA is 100 μM of each fatty acid conjugated to BSA (200 μM total). Without PA/OA conditions contain fatty acid free BSA. G) Isotopologue enrichment of palmitate measured by GC-MS. Cells were cultured in DMEM + 10% D2O + 10% dFBS +100 μM acetate for 24 hours. Bars represent mean of 3 or 4 biological replicates for each cell line. Statistical analysis performed on total hydrogen enrichment. Statistical significance was calculated by one-way ANOVA. CDT, charcoal dextran treated. Each point represents a biological replicate and error bars represent standard deviation. *p≤0.05; **p≤0.01; ***p≤0.001; ****p≤0.0001

To begin to investigate functional changes in lipid metabolism, we first tested if loss of ACLY and ACSS2 resulted in dependence on exogenous lipids. For this, DKO cells were cultured in media supplemented with serum treated with charcoal-dextran (CDT), a process that removes lipophilic compounds. DKO cells cultured in these lipid-depleted conditions failed to proliferate and began to undergo apoptosis (Figure 3D, E, F), while their ACLY KO counterparts proliferated with only a slight defect (Figure 3D). Interestingly, DKO cells also failed to proliferate in dialyzed serum (dFBS), which removes small molecules using a 10,000 MW cutoff membrane (Supplemental Figure 3G). Acetate increased proliferation in ACLY KO cells but had no effect on DKO cells (Figure 3D; Supplemental Figure 3E). Addition of BSA-conjugated palmitic and oleic acids (PA/OA) fully rescued proliferation of DKO cells, and either PA or OA alone was also sufficient (Figure 3D, E; Supplemental Figure 3E, F, G). Supplementation of mevalonate, mevalonate-phosphate, or the medium chain fatty acid octanoate failed to rescue DKO cell proliferation (Figure 3D; Supplemental Figure 3E). Thus, cells lacking ACLY and ACSS2 are dependent on exogenous fatty acids for viability and proliferation.

We next asked if this requirement for exogenous fatty acids reflected a limited ability to synthesize fatty acids *de novo*. Since the pathways and carbon sources supplying acetyl-CoA in the DKO cells were unknown, DKO cells were cultured in the presence of deuterated water and deuterium incorporation into palmitate was used to examine total *de novo* lipogenesis. Compared to WT and ACSS2 KO cells, DKO cells exhibited very low palmitate labeling (Figure 3G, Supplemental Figure 3H). This limited *de novo* lipogenesis suggests that DKO cells rely on exogenous fatty acids for proliferation due to reduced ability to adequately synthesize their own fat despite maintaining higher or similar concentrations of malonyl-CoA (Figure 2C).

### Fatty acids regulate histone acetylation in the absence of ACLY and ACSS2

To further understand how acetyl-CoA is used in the DKO cells, we assessed levels of histone acetylation, another major nutrient-sensitive acetyl-CoA-dependent process, across the 4 genotypes. We quantified acetylation at sites on histone H3 by mass spectrometry, focusing on two high abundance acetylation sites that have been proposed as acetate reservoirs, H3K23ac and H3K14ac^23^. Consistent with prior studies^4,24^, ACLY KO cells maintain lower levels of histone acetylation at H3K23 and H3K14 than their WT and ACSS2 KO counterparts, and DKO cells exhibit similar levels of acetylation as single ACLY KO cells (Figure 4A, B). To investigate if the capacity of DKO cells to acetylate is intact, we blocked histone deacetylation with the broad HDAC inhibitor trichostatin A (TSA). Here we analyzed the putative reservoir site H3K23ac, the regulatory site H3K27ac, and pan acetyl-H4. TSA treatment causes an increase in global histone acetylation in all marks analyzed over time in ACLY KO cells, and this effect is comparable in DKO cells (Figure 4C). This suggests that acetyl-CoA is readily available for use for histone acetylation in cells lacking both ACLY and ACSS2. Similarly, tubulin acetylation and total lysine acetylation dynamics, as assessed by a pan K-ac antibody, were comparable between genotypes (Figure 4D; Supplemental Figure 4A), indicating that acetyl-CoA is available in both the nucleus and cytosol in these cells.

**Figure 4:**
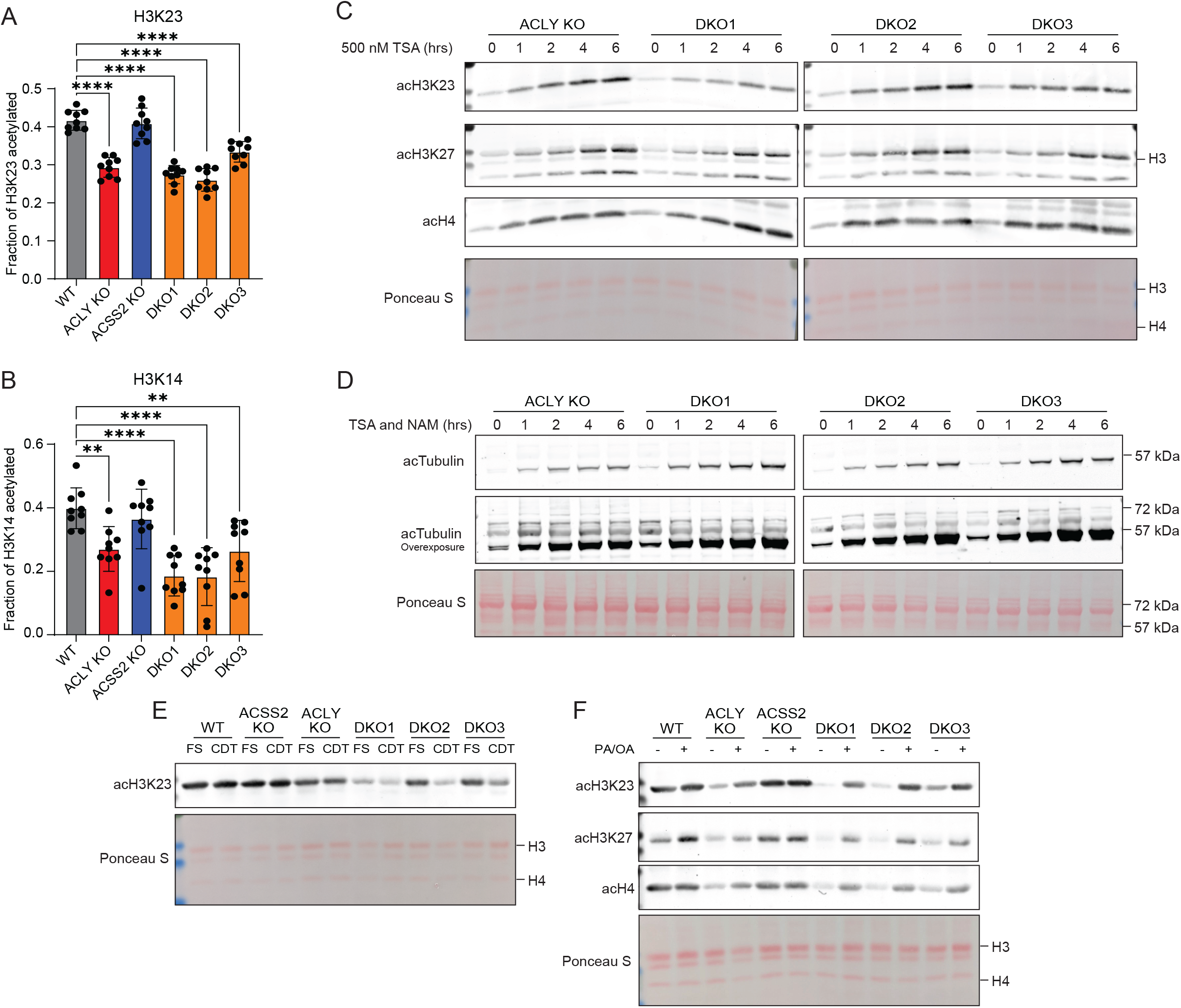
Fatty acid availability modulates histone acetylation independent of ACLY and ACSS2. A) Fraction of the total quantified H3K23 residues acetylated by LC-MS. Statistical significance was calculated by one-way ANOVA. B) Fraction of the total quantified H3K14 residues acetylated by LC-MS. Statistical significance was calculated by one-way ANOVA. C) Acid extracted histone western blot from cells grown in DMEM + 10% FS and treated with 500 nM trichostatin A (TSA) over a time course. Ponceau S stain for total protein in histone extracts used for western blot. D) Whole cell protein extract western blot from cells grown in DMEM + 10% FS and treated with 500 nM TSA and 500 μM nicotinamide (NAM) over a time course. Ponceau S stain for total protein in histone extracts used for western blot. E) Acid extracted histone western blot from cells cultured in DMEM + 10% FS or CDT for 24 hours. Ponceau S stain for total protein in histone extracts used for western blot. F) Acid extracted histone western blot from cells cultured for 24 hours in DMEM + 10% CDT and supplemented with PA/OA. PA/OA is 100 μM of each fatty acid conjugated to BSA (200 μM total). Ponceau S stain for total protein in histone extracts used for western blot. Each point represents a biological replicate and error bars represent standard deviation. *p≤0.05; **p≤0.01; ***p≤0.001; ****p≤0.0001

We hypothesized that serum lipids might play a role in sustaining histone acetylation independent of ACLY and ACSS2, because fatty acids oxidation produces acetyl-CoA and genes involved in oxidation are upregulated in DKO cells. To test this, DKO cells were incubated in lipid-depleted culture conditions. Lipid depletion led to a dramatic depletion of histone acetylation within 24 hours, and this was rescued by addition of PA/OA (Figure 4E, F; Supplemental Figure 4B). Acetyl-CoA abundance increased modestly with PA/OA supplementation, with some variability between lines (Supplemental Figure 4C). Malonyl-CoA, on the other hand, was potently suppressed by fatty acid supplementation (Supplemental Figure 4D). This is consistent with the known role of fatty acyl-CoAs in allosteric inhibition of the acetyl-CoA consuming enzyme ACC ^25,26^, which may divert acetyl-CoA towards histone acetylation, as previously reported ^27–29^. Consistently, ACC inhibition (ND630) also modestly increased histone acetylation and proliferation in the DKO cells while suppressing malonyl-CoA levels (Supplemental Figure 4D, E, F), indicating that fatty acids may act to increase histone acetylation in part through acetyl-CoA sparing from ACC consumption. We also tested octanoate, which has been previously shown to promote histone acetylation ^30^ and found that octanoate also increased histone acetylation and acetyl-CoA abundance in the DKO cells, though did not rescue proliferation (Supplemental Figure 4G, H; Supplemental Figure 3A). Together, these data show that exogenous fatty acids can regulate histone acetylation levels independently of ACLY and ACSS2.

### Fatty acids and glucose can feed acetyl-CoA pools and histone acetylation independent of ACLY and ACSS2

To determine if fatty acid oxidation contributes substantially to the acetyl-CoA pool in DKO cells, we used stable isotope tracing of uniformly labeled ^13^C palmitate (^13^C_16_-PA) into whole cell acetyl-CoA pools (Figure 5A). A time-course experiment showed maximal labeling of acetyl-CoA in DKO cells within 2 hours, reaching up to 30% (Supplemental Figure 5A). These findings suggest rapid and substantial breakdown of palmitate into acetyl-CoA. We also examined labeling from ^13^C-Glc, unexpectedly finding that although glucose was a minor contributor to acetyl-CoA pools in ACLY KO cells, it labeled a much greater fraction, similar to that of palmitate, in DKO cells (Figure 5B). Under these conditions, whole cell acetyl-CoA abundance was similar between genotypes (Supplemental Figure 5B). Additionally, in the absence of fatty acids, glucose labeled nearly 60% of the acetyl-CoA pool in DKO cells, though the pool size was reduced (Figure 5C; Supplemental Figure 5C). Malonyl-CoA accumulates in the absence of fatty acids in these cells in an ACC-dependent manner (Supplemental Figure 4D), and labeling paralleled that of acetyl-CoA, suggesting that glucose-derived carbon may feed into an extramitochondrial acetyl-CoA pool in these cells (Figure 5C). Acetate did not detectably contribute to acetyl-CoA pools in DKO cells (Figure 5C). Thus, both glucose and fatty acids are major carbon sources feeding acetyl-CoA pools in the absence of ACLY and ACSS2.

**Figure 5:**
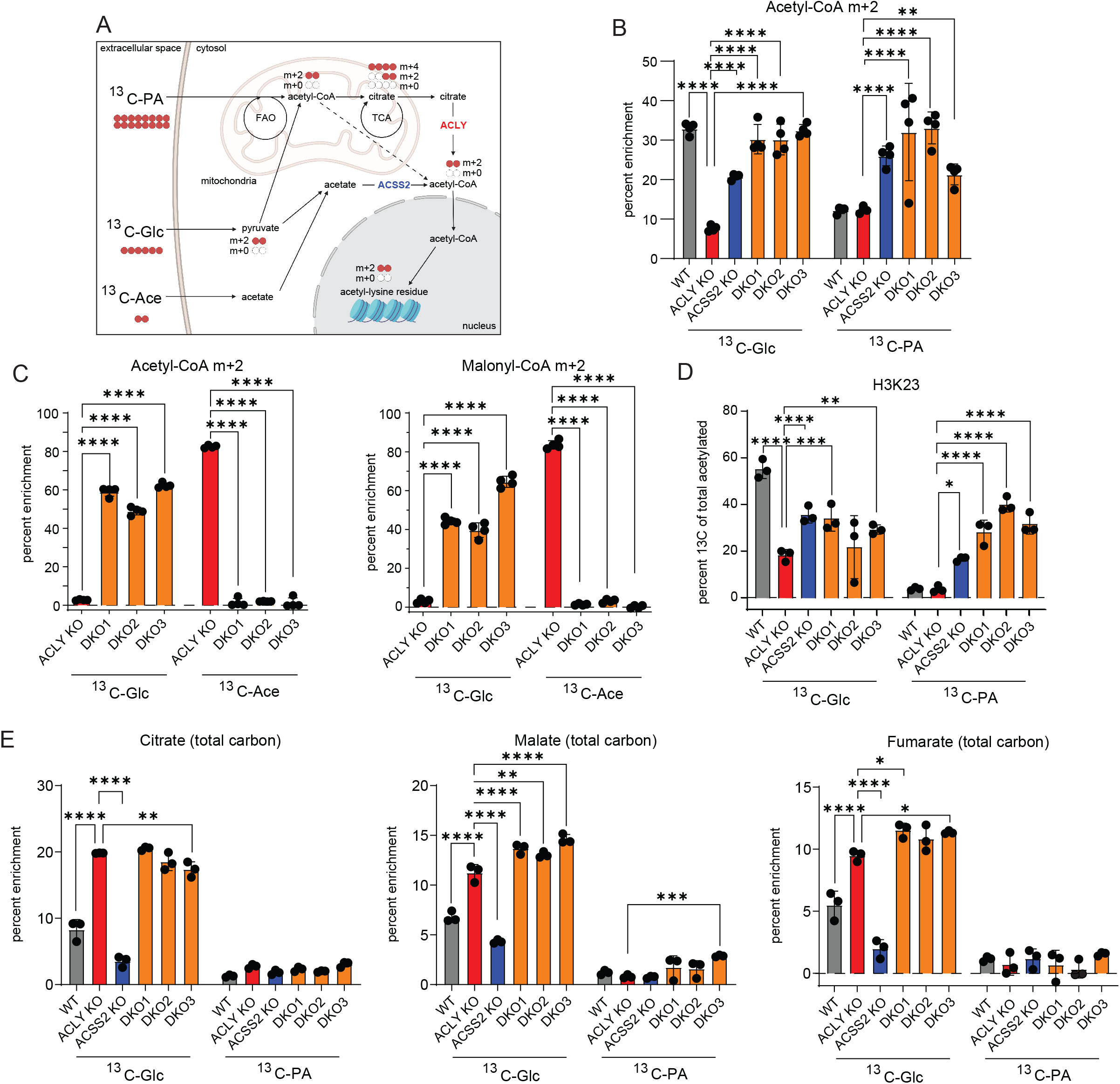
Fatty acids and glucose can supply acetyl-CoA for histone acetylation in a manner independent of ACLY and ACSS2. A) Schematic depicting glucose, palmitate, and acetate carbon tracing into acetyl-CoA and histone acetylation. Created with BioRender.com. B) ^13^C_6_-glucose and ^13^C_16_-palmitate tracing into acetyl-CoA analyzed by LC-MS. Cells were cultured in glucose and glutamine free DMEM + 10% CDT supplemented with 4 mM glutamine and either 10 mM ^13^C_6_-glucose and 100 μM palmitate conjugated to BSA or 10 mM glucose and 100 μM ^13^C_16_-palmitate conjugated to BSA for 2 hours. Statistical significance was calculated by two-way ANOVA. C) Acetyl-CoA and malonyl-CoA enrichment from ^13^C_6_-glucose or ^13^C_2_-acetate. Cells were cultured in glucose and glutamine free DMEM + 10% dFBS supplemented with 4 mM glutamine and either 10 mM ^13^C_16_-glucose and 100 μM acetate or 10 mM glucose and 100 μM ^13^C_2_-acetate for 6 hours. Statistical significance was calculated by two-way ANOVA. D) ^13^C_6_-glucose and ^13^C_6_-palmitate tracing into acetylation on extracted histones, analyzed by LC-MS. Cells were cultured in glucose and glutamine free DMEM + 10% CDT supplemented with 4 mM glutamine and either 10 mM ^13^C_6_-glucose and 100 μM palmitate conjugated to BSA or 10 mM glucose and 100 μM ^13^C_16_-palmitate conjugated to BSA for 24 hours. Statistical significance was calculated by two-way ANOVA. E) ^13^C_6_-glucose and ^13^C_16_-palmitate tracing into TCA cycle intermediates, analyzed by GC-MS. Cells were cultured in glucose and glutamine free DMEM + 10% CDT supplemented with 4 mM glutamine and either 10 mM ^13^C_16_-glucose and 100 μM palmitate conjugated to BSA or 10 mM glucose and 100 μM ^13^C_16_-palmitate conjugated to BSA for 6 hours. Statistical significance was calculated by two-way ANOVA. Each point represents a biological replicate and error bars represent standard deviation. All tests compared to ACLY KO as the control. *p≤0.05; **p≤0.01; ***p≤0.001; ****p≤0.0001

In order to determine if acetyl-CoA generated from glucose and palmitate is used for acetylation in the nucleus, we performed LC-MS analysis of histone acetylation in cells incubated with isotope labeled glucose or palmitate. WT cells used glucose derived carbons for histone acetylation, and this was blunted by ACLY KO as expected (Figure 5D). ACSS2 KO slightly blunted glucose carbon incorporation into histone acetylation, possibly reflecting the recycling of acetyl-groups over the 24-hour time^11^. DKO cells had greater histone acetylation labeling from glucose than ACLY KO, in line with acetyl-CoA labeling. Additionally, palmitate-derived carbon was used prominently by ACSS2 KO and DKO cells. The cumulative fractional labeling from palmitate and glucose in DKO cell histone acetylation matches that by WT and ACSS2 KO cells, while ACLY KO cells have much lower cumulative labeling from these two sources, consistent with unlabeled acetate feeding the acetyl-CoA pool only in those cells (Supplemental Figure 5D). Together, these data indicate that both glucose and fatty acids can supply nuclear-cytosolic acetyl-CoA through a pathway that does not require ACLY or ACSS2 and that can be used for histone acetylation.

### ACLY KO cells have altered glucose usage in the TCA cycle

We next sought to define the mechanism(s) of ACLY- and ACSS2-independent production of nuclear-cytosolic acetyl-CoA. Since both glucose and fatty acids could contribute to histone acetylation, we reasoned that this could occur either by two distinct substrate-specific mechanisms or a mechanism in which these substrates can feed into a common precursor acetyl-CoA pool (e.g., in mitochondria) that is then transported to the nuclear-cytosolic compartment (Supplemental Figure 5E).

Given multiple publications documenting a nuclear PDC^15–18^, we investigated the localization of PDH in the HCC cells. Using immunofluorescence and confocal microscopy, we observed prominent mitochondrial but minimal nuclear localization of catalytic subunit of the pyruvate dehydrogenase complex (PDHe1α) (Supplemental Figure 6A). Additionally, PDHe1α protein levels are unchanged in standard culture conditions and are unaffected by fatty acid availability (Supplemental Figure 6B, C). While not formally ruling out a role for a nuclear PDC, the data suggest that a different mechanism likely sustains the DKO cells.

We next asked if a mitochondrial route might be involved since both glucose and fatty acids can feed mitochondrial acetyl-CoA pools (Figure 5A; Supplemental Figure 5E). ^13^C-glucose labeling into TCA cycle intermediates was elevated in ACLY KO and DKO cells despite modest to no change in abundance of these metabolites (Figure 5E, Supplemental Figure 5F). Palmitate was poorly used as a carbon source in the TCA cycle in all genotypes, suggesting that fatty acid oxidation in the mitochondria is a more minor source of mitochondrial acetyl-CoA (Figure 5E). Inhibition of mitochondrial (etomoxir) and peroxisomal (thioridazine) fatty acid oxidation revealed that each organelle contributes to approximately half of acetyl-CoA produced from palmitate in the DKO cells (Supplemental Figure 5G). Together, the data indicate that glucose derived carbon feeds the mitochondrial metabolite pool and suggest the mitochondria as a possible intermediate location for glucose carbons before supplying nuclear-cytosolic acetyl-CoA in DKO cells.

### The carnitine shuttle facilitates glucose-dependent lipogenesis independent of ACLY

Acetyl-CoA cannot directly cross organelle membranes, and thus a transport mechanism out of mitochondria and/or peroxisomes is needed to explain the data. We noticed that *Crat*, encoding the carnitine acetyltransferase (CrAT) is upregulated in DKO cells (Fig. 3C). CrAT is present in mitochondria and peroxisomes ^31,32^ and transfers the acetyl moiety from acetyl-CoA onto carnitine to generate acetylcarnitine for organelle export via the carnitine-acylcarnitine translocase (CACT; SLC25A20). CrAT is thought to play a buffering function to prevent high levels of mitochondrial acetyl-CoA, which can suppress pyruvate oxidation and cause non-enzymatic acetylation^33–35^. CrAT is a reversible enzyme, and thus if CrAT is also present within the cytosol or nucleus, it could enable the regeneration of acetyl-CoA in the cytosol or nucleus from acetylcarnitine. Some evidence suggests it may be present and functional in the cytosol and nucleus, although such a role remains poorly understood ^20,36–38^.

To determine if acetylcarnitine could serve as a metabolic intermediate in the generation of acetyl-CoA in the nuclear-cytosolic compartment in DKO cells, we traced palmitate and glucose into acetylcarnitine. Acetylcarnitine is highly labeled from glucose in ACLY KO and DKO (Figure 6A), similar to TCA cycle labeling, suggesting that acetylcarnitine reflects and may be in equilibrium with the mitochondrial acetyl-CoA pool. Palmitate labels the acetylcarnitine pool, but to a lesser extent than glucose (Figure 6A). These findings prompted us to ask if CrAT provides a mechanism for shuttling of acetyl-units, especially those derived from glucose, into the nuclear-cytosolic acetyl-CoA pool.

**Figure 6:**
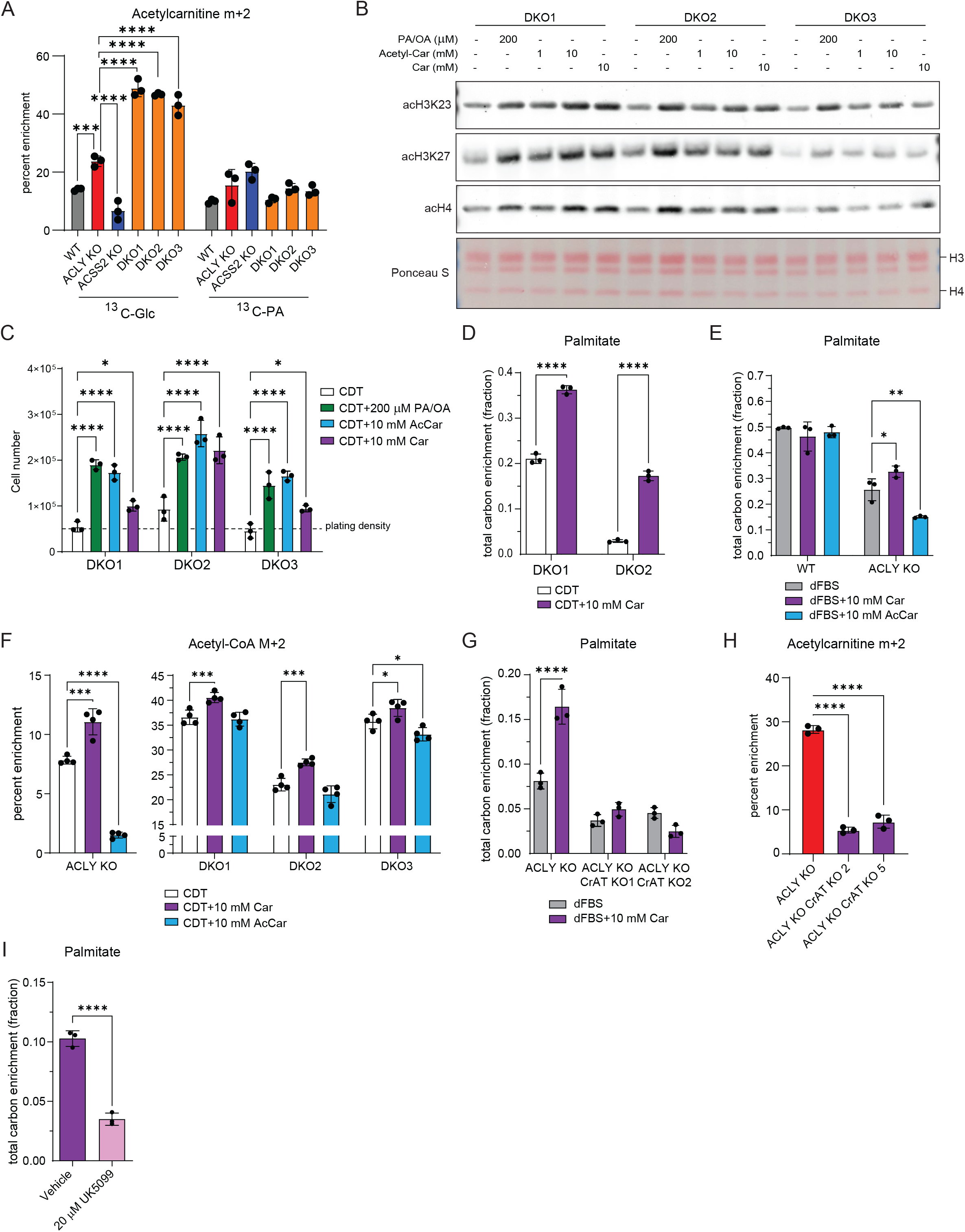
Carnitine facilitates histone acetylation and de novo lipogenesis from glucose derived carbon. A) ^13^C_6_-glucose and ^13^C_16_-palmitate tracing into acetylcarnitine, analyzed by LC-MS. Cells were cultured in glucose and glutamine free DMEM + 10% CDT supplemented with 4 mM glutamine and either 10 mM ^13^C_6_-glucose and 100 μM palmitate conjugated to BSA or 10 mM glucose and 100 μM ^13^C_16_-palmitate conjugated to BSA for 6 hours. Statistical significance was calculated by two-way ANOVA. B) Acid extracted histone western blot from cells cultured in DMEM + 10% CDT for 24 hours supplemented with PA/OA, acetylcarnitine, or carnitine. PA/OA is 100 μM of each fatty acid conjugated to BSA (200 μM total). Ponceau S stain for total protein in histone extracts used for western blot. C) Cell proliferation after 96 hours. Cells were plated in DMEM/F12 media overnight then cultured in DMEM + 10% CDT serum with or without the addition of metabolites. PA/OA is 100 μM of each fatty acid conjugated to BSA (200 μM total). Statistical significance was calculated by two-way ANOVA. D) ^13^C_6_-glucose tracing into palmitate measured by GC-MS. Cells were cultured in glucose and glutamine free DMEM + 10% CDT supplemented with 4 mM glutamine and 10 mM ^13^C_6_-glucose with or without 10 mM carnitine for 48 hours. Statistical significance was calculated by unpaired t-tests. E) ^13^C_6_-glucose tracing into palmitate measured by GC-MS. Cells were cultured in glucose and glutamine free DMEM + 10% dFBS supplemented with 4 mM glutamine and 10 mM ^13^C_6_-glucose with or without 10 mM carnitine or 10 mM acetylcarnitine for 48 hours. Statistical significance was calculated by two-way ANOVA. F) ^13^C_6_-glucose tracing into acetyl-CoA, analyzed by LC-MS. Cells were cultured in glucose and glutamine free DMEM + 10% dFBS supplemented with 4 mM glutamine and 10 mM ^13^C_6_-glucose supplemented with or without 10 mM carnitine or 10 mM acetylcarnitine for 6 hours. Statistical significance was calculated by two-way ANOVA. G) ^13^C_6_-glucose tracing into palmitate measured by GC-MS. Cells were cultured in glucose and glutamine free DMEM + 10% dFBS supplemented with 4 mM glutamine and 10 mM ^13^C_6_-glucose with or without 10 mM carnitine for 48 hours. Statistical significance was calculated by unpaired t-tests. H) ^13^C_6_-glucose tracing into acetylcarnitine, analyzed by LC-MS. Cells were cultured in glucose and glutamine free DMEM + 10% dFBS supplemented with 4 mM glutamine and 10 mM ^13^C_6_-glucose for 6 hours. Statistical significance was calculated by one-way ANOVA.I) ^13^C_6_-glucose tracing into palmitate measured by GC-MS. Cells were cultured in glucose and glutamine free DMEM + 10% dFBS supplemented with 4 mM glutamine and 10 mM ^13^C_6_-glucose with 10 mM carnitine with vehicle control or 20 μM UK5099 for 48 hours. Statistical significance was calculated by unpaired t-tests. Each point represents a biological replicate and error bars represent standard deviation. *p≤0.05; **p≤0.01; ***p≤0.001; ****p≤0.0001

If CrAT can generate acetyl-CoA for histone acetylation, we anticipated that the supplementation of acetylcarnitine to the culture media should boost histone acetylation and proliferation in lipid-depleted DKO cells, and this was indeed the case (Figure 6B, C). Notably, supplementation of DKO cells with L-carnitine alone was also able to increase both histone acetylation and cell proliferation in lipid depleted conditions (Figure 6B, C), suggesting that it may promote acetylcarnitine shuttling out of mitochondria.

To further probe this possibility, we tested if L-carnitine supplementation increased the ability of DKO cells to carry out de novo lipogenesis. Using deuterated water labeling, we found that L-carnitine significantly increased synthesis of palmitate and stearate in DKO cells (Supplemental Figure 7A). Additionally, supplementation of L-carnitine enhanced glucose dependent de novo lipogenesis in the absence of ACLY and ACSS2, implicating the carnitine shuttle in transport of acetyl-units out of mitochondria (Figure 6D; Supplemental Figure 6B).

To understand if carnitine can also promote glucose dependent *de novo* lipogenesis outside of the DKO context, we tested the impact of carnitine supplementation on WT and ACLY KO cells. WT cell *de novo* lipogenesis from glucose was unaffected by carnitine supplementation (Figure 6E; Supplemental Figure 7C, D, E, F). ACLY KO cells, however, showed a significant increase in glucose derived *de novo* lipogenesis when supplemented with carnitine (Figure 6E; Supplemental Figure 7C, D, E, F). Additionally, supplementation of acetylcarnitine further reduced glucose-dependent *de novo* lipogenesis in ACLY KO cells, suggesting that the acetyl-units provided by acetylcarnitine supplementation dilute the glucose-derived acetyl-CoA pools (Figure 6E; Supplemental Figure 7C, D, E). Supporting this model, glucose incorporation into whole cell acetyl-CoA was increased in ACLY KO and DKO cells with carnitine supplementation, and acetylcarnitine suppressed glucose-dependent labeling, particularly in ACLY KO cells (Figure 6F). Together, these findings show that acetyl-units from the mitochondria can be exported and used in the cytosol independent of ACLY in a manner facilitated by carnitine.

To test if CrAT is necessary for the carnitine-mediated shuttling of acetyl-units out of the mitochondria, we generated ACLY KO/CrAT KO cells (Supplemental Figure 7G, H). CrAT deficiency suppressed the residual glucose dependent *de novo* lipogenesis in ACLY KO cells and abrogated the effect of carnitine supplementation in promoting *de novo* lipogenesis (Figure 6G, Supplemental Figure 7I). Additionally, CrAT KO suppressed acetyl-carnitine labeling from glucose (Figure 7H). Next, we tested if the mitochondrial entry of pyruvate was necessary for carnitine driven glucose labeling of fatty acids in ACLY KO cells by inhibiting the mitochondrial pyruvate carrier (MPC). Inhibition of MPC by UK5099 causes a dramatic decrease in glucose contribution to *de novo* lipogenesis (Figure 6I, Supplemental Figure 7J). Notably, UK5099 in combination with carnitine supplementation appeared to cause toxicity (Supplemental Figure 7K). These data indicate that the mitochondrial carnitine shuttle and CrAT facilitate an alternative pathway for two carbon units to leave mitochondria for use in nuclear-cytosolic processes.

**Figure 7:**
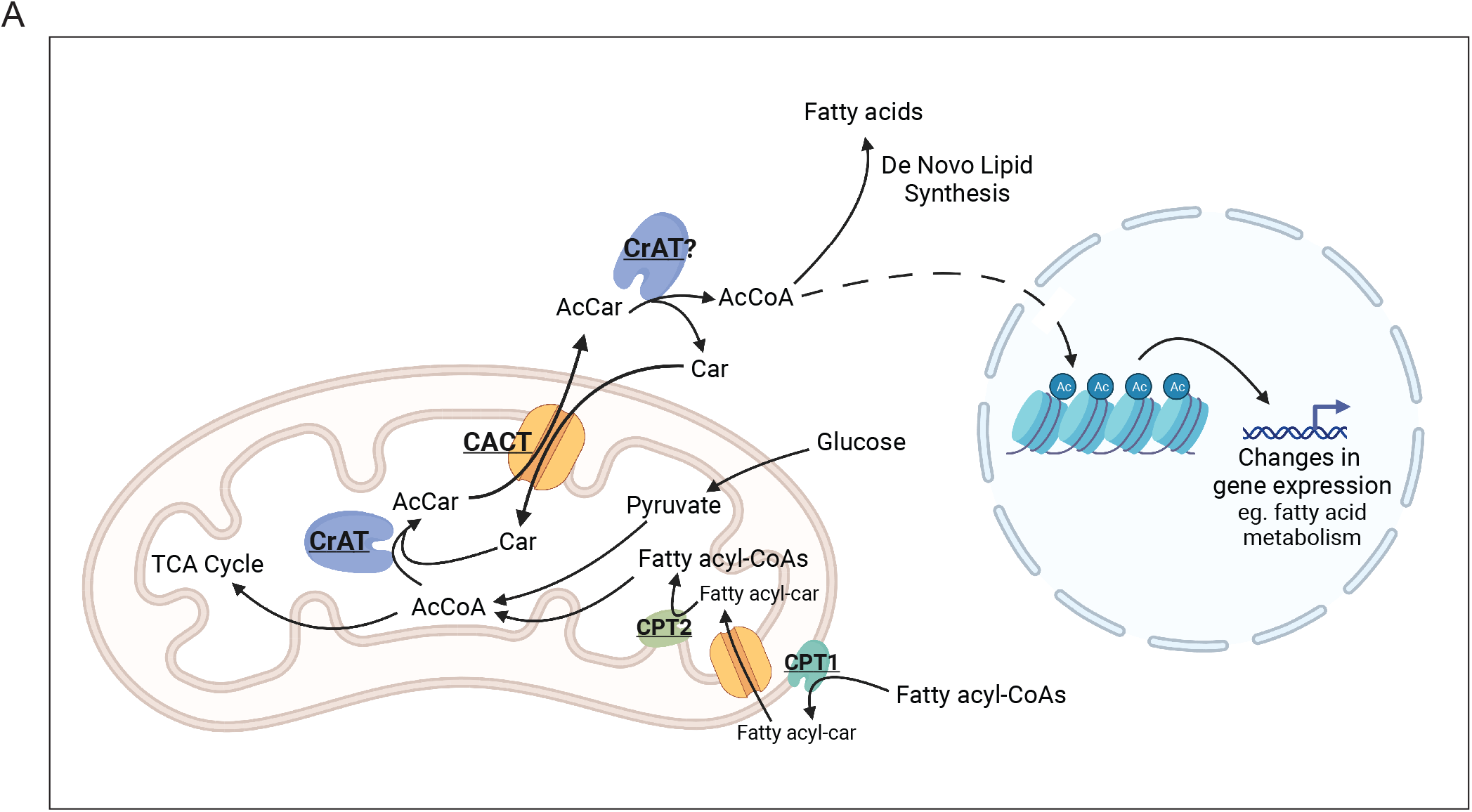
The carnitine shuttle provides acetyl-units to the nuclear cytosolic compartment. A) Schematic depicting CrAT-dependent acetylcarnitine shuttling out of the mitochondria for acetyl-CoA generation in the nuclear-cytosolic compartment. Arrows represent biochemical conversions. Created with BioRender.com.

## DISCUSSION

The two well established routes to nuclear-cytosolic acetyl-CoA pools used for lipid synthesis and histone acetylation are via mitochondrial citrate export and cleavage by ACLY and acetate activation by ACSS2. In this study we demonstrate that the carnitine shuttle and CrAT can provide an alternative route for two carbon transport from mitochondria to cytosol. In cancer cell lines lacking both ACLY and ACSS2, we show that a pool of cytosolic acetyl-CoA is maintained, that histone acetylation is active, and that both glucose and fatty acids can supply acetyl-CoA. We further show that fatty acids boost histone acetylation both by serving as a carbon source and via acetyl-CoA sparing by ACC inhibition. In the absence of ACLY, we show that carnitine supplementation increases *de novo* lipogenesis from glucose in a CrAT-dependent manner, thus demonstrating that the carnitine-CrAT pathway can be used as an ACLY-independent means of transporting two carbon units to the cytosol (Figure 7A). Of note, our tracing data also suggest that a peroxisomal route is likely important for at least a portion of fatty acid-dependent acetyl-CoA production. This is confirmed and studied in depth in a complementary study reported in a manuscript co-submitted with this one (Kumar et al, co-submitted). Overall, this work broadens the understanding of nuclear-cytosolic acetyl-CoA metabolism and opens new avenues for investigation into the regulation of de novo lipogenesis and histone acetylation.

One reason for undertaking this study to identify compensatory mechanisms of acetyl-CoA metabolism is that both ACLY and ACSS2 are of interest as therapeutic targets. The liver specific ACLY inhibitor bempedoic acid is FDA approved for LDL cholesterol lowering^3,39^, and the first ACSS2 inhibitor has entered oncology clinical trials (NCT04990739). At least in mice, ACLY deletion^6^ or bempedoic acid^40^ strongly suppresses glucose-dependent hepatic *de novo* lipogenesis, suggesting that ACLY is a dominant mechanism for glucose-dependent *de novo* lipogenesis in the liver, although acetate can sustain lipogenesis in the absence of ACLY^6^. However, looking to the future, ACLY is also of interest as a potential target in oncology, and given the metabolic plasticity of cancer cells and the ability to shift between nutrient sources^41^, a potential role of CrAT should be considered. Carnitine is synthesized in the liver, but also can be obtained through the diet. Carnitine is abundant in certain diets, such as those high in red meat, and the impact of dietary carnitine warrants exploration. Additionally, L-carnitine and acetyl-L-carnitine dietary supplements are widely available. Inhibiting carnitine metabolism would need to be evaluated with caution, as carnitine deficiency can cause severe phenotypes including muscle wasting, heart failure, liver damage, and cognitive delays^42^. In addition, it will be of interest to understand whether the carnitine shuttle plays a role in compensating for ACLY and ACSS2 under conditions such as high fat diet, in which both enzymes are suppressed in adipose tissue^43^; presumably acetyl-CoA would still be needed for histone acetylation and the mevalonate pathway even if fatty acid synthesis activity is low.

CrAT has mainly been studied for its role in mitochondrial metabolism, including fatty acid oxidation and acetyl-CoA buffering ^33–35,44^; however there have been prior reports of CrAT activity outside of the mitochondria, as well as observations consistent with such activity. Early biochemical characterization of CrAT activity showed that it was at least partially localized on the cytosolic face of the ER membrane^36^. Yet, most reports suggest that CrAT is not a membrane bound protein and is localized within the mitochondrial lumen and the peroxisome, predominantly due to different CrAT isoform expression^32^. Tracing studies using labeled acetylcarnitine show that the acetyl-units from acetylcarnitine can indeed be used for lipid synthesis, and that their usage is increased with ACLY inhibition by hydroxycitrate^37^. Another study suggested that an isoform of CrAT is present in the nucleus and may promote histone acetylation in a manner dependent on the carnitine/acylcarnitine translocase (CACT)^20^. Finally, acetyl-proteomics data in CrAT KO skeletal muscle showed that this perturbation increases mitochondrial acetylation, supporting the idea that CrAT can act as a buffer system for mitochondria acetyl-CoA, but also observed decreases in cytosolic protein acetylation^33^. In addition to acetyl-CoA, CrAT may also be important for shuttling of other short-chain acyl-CoAs from mitochondria to the nuclear-cytosolic compartment. This is consistent with evidence that odd chain and branched chain fatty acid synthesis is CrAT dependent^38^, and that isoleucine catabolism, a mitochondrial process, can supply propionyl-CoA for histone propionylation and that this correlates with production of propionyl-carnitine^8^. Nevertheless, despite these reports in the literature, the significance of such a route has not been widely appreciated. Our findings, using genetic models and isotope tracing, are consistent with a model in which CrAT produces acetylcarnitine from acetyl-CoA in mitochondria and then following export to the cytosol, it converts acetylcarnitine back to acetyl-CoA as an alternative to ACLY for two carbon transfer from mitochondria (Figure 7A). However, while CrAT is required for this pathway in mitochondria, it remains possible that another unknown enzyme or a non-enzymatic process is responsible for nuclear-cytosolic conversion of acetylcarnitine into acetyl-CoA.

Further work will be needed to characterize the physiological contexts in which this pathway is employed versus the ACLY- or ACSS2-dependent routes to supply acetyl-CoA for lipid synthesis and chromatin modification. ACLY has been shown to participate in a non-canonical TCA cycle in a manner that influences cell fate^45^. This depends on oxaloacetate production by the ACLY reaction, which is returned to the TCA cycle after conversion to malate by MDH1. The carnitine shuttle offers a means to transport two carbon units to the cytosol without concomitant oxaloacetate production and thus one prediction is that this route might be important for supporting histone acetylation or lipogenesis in cell types that do not engage the non-canonical cycle.

An additional question emerging from this work is the function of histone acetate reservoirs. It has been proposed that histones provide a large reservoir of acetyl-units that can be mobilized either under metabolic stress or for a ready source of acetyl-CoA for site-specific histone acetylation and gene regulation^23,46,47^. Lipid deprivation in the DKO cells causes a rapid depletion in global histone acetylation in DKO cells. This model could uniquely enable studies of the consequences of histone acetylation reservoir depletion for gene regulation and chromatin structure.

Overall, this work provides evidence that the carnitine shuttle can contribute to both lipid synthesis and metabolites needed for chromatin modification. These data lay the groundwork for functional studies into the role of this pathway in physiological and disease contexts in regulation of cell state via histone acetylation and viability and growth potential via lipid synthesis.

## MATERIALS AND METHODS

### Cell lines

Murine Aclyf/f hepatocellular carcinoma cell lines were generated from diethylnitrosamine induced tumors in Aclyf/f mice and have been previously described^8^. ACLY KO cells were generated by administration of adenoviral Cre recombinase, and single cell clonal populations were generated by limiting serial dilution. WT and ACLY KO cells were then transduced with a LentiCRISPR v2 vector with no guide RNA or containing a guide RNA targeting the first exon of ACSS2 or near the active site of CrAT:

ACSS2 KO, DKO1, DKO2 (and KPACs):
mACSS2sg2 - CGAGCTGCACCGGCGTTCTG
DKO3:
mACSS2sg6 - CTGCACCGGCGTTCTGTGG
ACLY KO CrAT KO1, ACLY KO CrAT KO2:
msgCRAT1F - CACCGTCCACAAGTGCAACTATGGG

Following transduction, cells were treated with puromycin until the entirety of an un-transduced cell population died. Following puromycin treatment single cell clonal populations were generated for ACSS2 KO, CrAT KO, ACLY/ACSS2 DKO, and ACLY/CrAT DKO cells by limiting serial dilution. The bulk cell of population of WT and ACLY KO cells transduced with the empty LentiCRISPR v2 were used as controls.

Murine pancreatic cancer cell lines were generated from pancreatic ductal adenocarcinoma tumors of Pdx1-Cre; LSL-KrasG12D; Tp53f/f; Aclyf/f mice^22^. A female mouse with palpable tumors in the peritoneal cavity was sacrificed at approximately 15 weeks of age. Pancreatic tumor was excised from the animal, minced in smaller pieces using sterile scissors and finally digested using a collagenase VI solution (2 mg/mL in DMEM/F12; Sigma #C9891) for 20 minutes at 37°C. The solution was then filtered through a 70 μM mesh to obtain a single cell suspension. Cells were cultured in PDEC medium^22^ and tested for mycoplasma contamination. Cells were passaged at confluency and ACLY deletion was confirmed by western blotting after 3 passages. Pancreatic cancer ACLY KO cells were then transduced with a retroviral pOZ-N vector (addgene 3781) expressing ACLY or an empty pOZ-N vector to generate cells with reconstituted ACLY (ACLY KO plus ACLY cDNA) or an empty vector (ACLY KO). Cells were selected using IL-2R selection beads via pull down. To generate ACSS2 KO cells, cells were then transduced with a LentiCRISPR v2 empty vector or vector containing a guide RNA targeting the first exon of ACSS2. Following transduction, cells were treated with puromycin until the entirety of an un-transduced cell population died. Following puromycin treatment single cell clonal populations were generated for ACLY KO cells with ACLY cDNA and ACSS2 KO as well as ACLY KO cells with ACSS2 KO by limiting serial dilution.

### Cell culture

Murine HCC cells were cultured in DMEM/F12 (Gibco #11320033) supplemented with 10% super calf serum (FS) (Gemini #100-510). Murine pancreatic cancer cells were cultured in DMEM (Gibco #11965084) supplemented with 10% super calf serum (FS) (Gemini #100-510). Cell growth experiments were performed by plating cells at the indicated density in DMEM/F12 + 10% FS and cells were allowed to adhere overnight. Culture medium was changed the following day to the indicated conditions. Media for all experiments used high glucose DMEM (Gibco #11965084) or glucose and glutamine free DMEM (Gibco #A1443001) supplemented with 10 mM glucose and 4 mM glutamine unless otherwise indicated. Dialyzed fetal bovine serum (dFBS) (Gemini #100-108) and charcoal stripped FBS (charcoal dextran treated - CDT) (Corning #35-072-CV) or charcoal stripped FBS (CDT) (Sigma F6765) were used when indicated. All cell lines were routinely tested for mycoplasma contamination.

Metabolite addback and inhibitor treatment experiments were performed in the media conditions listed with supplementation of the listed metabolite or drug with equal volumes of vehicle control unless otherwise specified. Acetate, pyruvate, and mevalonate were supplemented after dissolving in water. Carnitine and acetylcarnitine were dissolved in media used for each experiment. Fatty acid supplementation used palmitate and/or oleate conjugated to fatty acid free BSA (Bioworld 22070023) and fatty acid free BSA in an equivalent volume was supplemented as control.

### Soft Agar Colony Formation Assay

Cells were plated in 6 well plates at a density of 2.5×10^4^ cells per well. First, plates were coated with glucose and glutamine free DMEM media containing 10 mM glucose and 4 mM glutamine and supplemented with 10% FS and 0.6% Bacto Agar. Cells were plated on top of the 0.6% Bacto agar layer in glucose and glutamine free DMEM media containing 10 mM glucose and 4 mM glutamine and supplemented with 10% FS and 0.3% Bacto Agar. Each cell line was plated in triplicate. Fresh media was added to the wells every 7 days for 3 weeks. Images were taken for analysis after 3 weeks. A total of 4 non-overlapping images were taken of each well totaling 12 images for analysis per cell line. Colonies were counted after blinding of images.

### Acid Extraction of Histones

Acid extraction on isolated nuclei was performed as previously described^4^. Cells were lysed with NIB-250 buffer (15 mM Tris-HCl (pH 7.5), 60 mM KCl, 15 mM NaCl, 5 mM MgCl^2^, 1 mM CaCl^2^, 250 mM sucrose, 1 mM DTT, 10 mM sodium butyrate, protease inhibitors) with 0.1% NP-40 for 5 minutes on ice. Nuclei were pelleted from the cell lysate by centrifugation at 600g at 4°C for 5 minutes. Extracted nuclei were washed with NIB-250 twice. Extracted nuclei were resuspending in 0.4N H2SO4 and rotated for 4 hours or overnight at 4°C to extract histone proteins. The extracts were cleared by centrifugation at 11,000g at 4°C for 10 minutes. Clarified histone extracts were precipitated by adding 100% TCA to a final concentration of 20% TCA and the extracts were at 4°C overnight. Precipitated histones were centrifuged at 11,000g at 4°C for 10 minutes. Histones were washed with 1 mL acetone + 0.1% 12 N HCl once and 1 mL acetone twice. The histone pellet was air dried at room temperature then resuspended in glass distilled H_2_O. Resuspended histones were used for western blotting.

### Western blotting

Cell for whole cell protein lysates were collected by trypsin mediated release from tissue culture plates. Cells were spun at 8000 rpm for 5 min and were kept on ice. Pellets were washed once with PBS. The cell pellet was resuspended in 50-100 μL RIPA buffer [1% NP-40, 0.5% deoxycholate, 0.1% SDS, 150 mM NaCl, 50 mM Tris plus protease inhibitor cocktail (Sigma-Aldrich, P8340) and phosSTOP if phosphorylated proteins were being investigated (Sigma-Aldrich, 04906845001)]. Cell lysis was allowed to occur on ice for 10 minutes. Cells were sonicated with a Branson Sonifier for 10 pulses at 20% amplitude. Cell lysate was clarified by centrifugation at 15,000×g for 10 minutes at 4°C and supernatant was transferred to a new tube. Samples were stored at −80C until analysis. All blots were developed using a LI-COR Odyssey CLx system. Antibodies used in this study were: ACLY (Proteintech #15421-1-AP), ACSS2 (CST #3658), Beta-actin (CST #3700), Alpha-tubulin (CST #2144), CrAT (Cloud-Clone Corp #PAC400Mu01), PDHe1α (Santa Cruz sc-377092), Pan-acetyl-lysine (CST #9441), Acetyl-tubulin (CST #3971), Acetyl-H3K23 (CST #14932), Acetyl-H4 (Millipore 06-866), Acetyl-H3K27 (Abcam ab4729), Acetyl-H3K9 (Active Motif AB_2793569), Acetyl-H4K5 (Millipore 07-327).

### RT-qPCR

RNA was extracted from cells after trypsinization and pelleting by centrifugation at 8000xg for 5 minutes. Pellets were resuspended in 500 μL Trizol (Life Technologies). RNA was extracted following the Trizol manufacturer protocol. Then, cDNA was prepared using high-capacity RNA-to-cDNA master mix (Applied Biosystems, 4368814) according to kit instructions. cDNA was diluted 1:20 and amplified with PowerUp SYBR Green Master Mix (Applied Biosystems, A25778) using a ViiA-7 Real-Time PCR system (Applied Biosystems). Fold change in expression was calculated by the ΔΔCt method using actin as a control. Primer sequences listed below.

CrAT-mF TGGTCATCTACTCCAGCCCA
CrAT-mR AACTGGCAGCGTCTCATTGT
Actin-mF TGGTGGGAATGGGTCAGAA
Actin-mR TCTCCATGTCGTCCCAGTTG

### D_2_O and glucose labeling of fatty acids and FAME GC-MS

Cells were seeded at a density of 7.5×10^5^ cells per plate in DMEM/F12 containing 10% FS. For deuterium tracing, the following day the media was changed to DMEM media containing 10% FS or 10% dFBS supplemented with 100 □M acetate and 10% deuterium oxide (Sigma 151882). For glucose tracing, the following day the media was changed to glucose and glutamine free DMEM media containing 10% dFBS or 10% CDT with 4 mM glutamine and 10 mM ^13^C_6_ glucose. After 24- or 48-hours cells were washed with ice cold DPBS and trypsinized. The trypsin reaction was stopped using cold 10% fatty acid free BSA in DPBS to remove any exogenous fatty acids. The cells were then washed twice with ice cold DPBS and the cell pellet was frozen at − 80°C until extraction.

Lipids were extracted by resuspending the cell pellet in 2 ml ice cold methanol, followed by addition of 700 μL ice cold glass distilled water. 10 μL of 1 mM heptadecanoic acid in methanol was added to each sample as an internal standard. Cell suspensions were sonicated using a Branson Sonifier 250 at an output of 2.5 and a duty cycle of 20% for 15 pulses. Following sonication, 1 mL of ice-cold chloroform was added, and the suspension was mixed by vortexing. An additional 700 μL of ice-cold chloroform and 700 μL of ice-cold glass distilled water was added to the mixture and vortexed to mix. The suspension was centrifuged at 8000g for 10 minutes at 4°C. The chloroform fraction was transferred to a new tube and the original suspension was re-extracted with 700 μL ice cold chloroform and centrifuged at 8000g for 10 minutes at 4°C. The chloroform fraction from both extractions were pooled and 100 μL ice cold water was added and the sample was vortexed to mix. The sample was centrifuged at 8000g for 10 minutes at 4°C and the chloroform fraction was transferred to a new tube and dried under nitrogen at 40°C.

Lipids were derivatized to methyl-esters by first resuspending the dried lipid extracts in 2 mL methanol: toluene (80:20) containing butylated-hydroxy toluene (5 mg in 50 mL). Acetyl-chloride (2 μL) was added, and the samples were heated to 95°C for 1 hour. Following heating, 5 mL 6% potassium carbonate was added, and the samples were centrifuged at 8000 RFC for 10 minutes at 4°C. The toluene layer was transferred to a new tube and centrifuged at 10000 RPM for 5 minutes at room temperature. The toluene layer was transferred to a glass GC/MS vial with a volume reducing insert. Fatty acid methyl esters were analyzed by GC/MS on an Agilent GC/MS 7890A/5975A with a DB-5 column. Deuterium or carbon enrichment into palmitate was determined using Fluxfix^48^.

### Acyl-CoA analysis by LC-MS

For extraction of acyl-CoAs, culture medium was completely aspirated from cells in 6cm plates before adding 1ml ice-cold 10% trichloroacetic acid to plates. For quantification experiments internal standard was added containing [^13^C_3_^15^N_1_]-labeled acyl-CoAs generated in pan6-deficient yeast culture^49^. Plates were scraped to collect the cells. Samples were then sonicated for 10 × 0.5 s pulses to completely disrupt cellular membranes and incubated on ice to precipitate proteins. Protein was pelleted at 16,000g for 10 min at 4°C. Supernatant was collected and purified by solid-phase extraction using Oasis HLB 1cc (30 mg) SPE columns (Waters). Eluate was evaporated to dryness under nitrogen gas and re-suspended in 50 μL of 5% 5-sulfosalicylic acid (w/v) for injection. Samples were analyzed by an Ultimate 3000 autosampler coupled to a Thermo Q-Exactive Plus Instrument in positive electrospray ionization (ESI) mode as previously described^50^. For quantitation, a calibration curve was generated using commercially available standards and internal standards containing [^13^C_3_^15^N_1_]-labeled acyl-CoAs generated in pan6-deficient yeast culture^49^ were added to each sample. For enrichment analysis, isotopically labeled glucose (^13^C_6_ glucose), acetate (^13^C_2_ acetate), or palmitate (^13^C_16_ palmitate) was added to culture media and the enrichment into acyl-CoAs was determined using FluxFix based on samples treated with no isotope tracer^48^.

### Acyl-carnitine analysis by LC-MS

For extraction of acyl-carnitines, cells were washed with 5ml ice-cold 0.9% NaCl to remove extracellular metabolites and scraped on ice in 1ml −80°C 80% HPLC-grade methanol/20% HPLC-grade water. Extracts were collected in 1.5ml tubes and 10ng of d3-propionyl-L-carnitine internal standard (Cayman 26579) in 50μl 80% HPLC-grade methanol/20% HPLC-grade water was added to each sample. Samples were vortexed and incubated at −80°C for 30 min following centrifugation at 17,000g for 10 min at 4°C. Supernatants were transferred into a 96-well plate and dried under nitrogen gas at room temperature overnight. Dried metabolites were resuspended in 50μl 95% HPLC-grade water/ 5% HPLC-grade methanol using TOMTEC QUADRA 4. For each sample, 1μl was injected and analyzed using a Vanquish Duo UHPLC system coupled to a Thermo Q-exactive Plus Orbitrap Instrument in positive electrospray ionization mode in full scan mode from 150-1000 m/z. The HPLC system used a hydrophilic interaction chromatography (HILIC) analytical column (Ascentis Express 2.1 mm ×150 mm, 2.7μm). The column was kept at 30°C and the flow rate was 0.5ml/min. The mobile phase was solvent A (10 mM ammonium acetate and 0.2% formic acid in water) and solvent B (10 mM ammonium acetate and 0.2% formic acid in 95% HPLC-grade acetonitrile/5% HPLC-grade water). Elution gradients were run starting from 98% B to 86% B from 0-7 min; 86% B to 50% B from 7-7.3 min; 50% B to 10% B from 7.3-8.3; 10% B was held from 8.3-14.5 min, 10% B to 98% B from 14.5 to 14.510; 98% was held from 14.510-15 min and then the column was equilibrated while eluting on the other identical column. For each analyte and the internal standard, the peak corresponding to the [M+H]^+^ ion at 5ppm was integrated in Tracefinder 4.1 (Thermo Scientific). The enrichment of isotopically labelled glucose (^13^C_6_ glucose) or palmitate (^13^C_16_ palmitate) into acetyl-carnitine was determined using FluxFix based on samples treated with no isotope tracer^48^.

### TCA metabolite analysis by GC-MS

Polar metabolites were extracted from cells through addition of 1mL 80:20 methanol:water chilled to −80°C. Cells were collected by scraping after addition of norvaline as an internal standard, lysed by 3 rounds of freeze thawing, and insoluble material was removed through centrifugation at 12,000g at 4°C for 10 minutes. The pellet was used for protein quantification after resuspension in 2% SDS 0.1 mM Tris buffer. The supernatant containing polar metabolites was evaporated to dryness by SpeedVac. The dried pellet was stored at −80°C until derivatization. Samples were derivatized by addition of 30 □L of 5 mg/mL methoxyamine in pyridine and heated for 15 minutes at 70°C. A total of 70 □ L MTBSTFA was then added, and the samples were heated at 70°C for 1 hour. After derivatization, samples were centrifuged at 12,000 rpm for 5 minutes. The supernatant was transferred to a glass GC/MS vial with a volume reducing insert. Samples were analyzed by GC/MS on an Agilent GC/MS 7890A/5975A with a DB-5 column. 13-C enrichment was determined using Fluxfix based on samples treated with no isotope tracer^48^. Relative quantification was performed by normalizing the sum of the AUC of all isotopologues for each metabolite to the sum of the AUC of all isotopologues of norvaline within each sample followed by normalization to the average protein quantification within each experimental group.

### Histone acetylation UPLC-MS/MS analysis

Histones were extracted as described in “Acid Extraction of Histones”. Dry histone pellets were stored at −80°C until processing for LC/MS analysis. The unmodified lysines in dry histone were propionylated by propionic anhydride at pH 8 and 51°C for 1 hour followed by trypsin digestion at pH 8 and 37°C for overnight as in our previously published procedures^51^. A Waters Acquity H-class UPLC coupled with a Thermo TSQ Quantum Access triple quadrupole mass spectrometer was used to quantify the acetylated lysines on H3 tryptic peptides. The UPLC and MS/MS settings, solvent gradient and detailed mass transitions were reported previously^52^. Retention time and specific mass transitions were both used to identify individual acetylated and propionylated peaks. The ^13^C-labeled acetyl-lysine peptide peaks were identified by considering the shift in mass by +2. The resolved peaks were integrated using Xcalibur software (version 2.1, Thermo). Relative quantitative analysis was used to determine the amount of modification on individual lysines.

### Stable isotope labeling of essential nutrients in cell culture-subcellular fractionation *(SILEC-SF) acyl-CoA quantitation

SILEC-SF was performed as previously described^8^. In brief, WT HCC cells (D42) were used to generate SILEC internal standard by passaging of cells in ^15^N^13^C_3_-pantothenate (Vitamin B5) for at least 9 passages as previously described^8^. SILEC WT cells were mixed with DKO1 cells before fractionation. Mitochondria and cytosolic fractions were separated through differential centrifugation. Extracted acyl-CoAs were subjected to analysis by LC-MS as described above.

### RNA-Sequencing

RNA was extracted from HCC cells cultured in DMEM + 10% FS for 24 hours using a Qiagen RNeasy Plus Mini Kit (Qiagen #74134) with an additional on column DNA digestion using Qiagen TURBO DNase (Qiagen #AM2238) based on manufacturer protocols.

RNA-seq libraries were prepared with the NEBNext poly(A) Magnetic Isolation Module (NEB #E7490L) followed by the NEBNext Ultra Directional RNA library preparation kit for Illumina (NEB #E7420L) according to manufacturer’s protocol. Library quality was assessed using an Agilent BioAnalyzer 2100 and libraries were quantified with the Library Quant Kit for Illumina (NEB #E7630L). Libraries were then diluted to 1.8pM and sequenced on the NextSeq500 platform using 75-base-pair (bp) single-end reads. All RNA-seq read alignment was performed using Illumina RNA-seq alignment software (version 2.0.1). Briefly, reads were mapped to Mus musculus University of California Santa Cruz (UCSC) mouse GRCm38/mm10 reference genome with RNA STAR aligner under default settings (version 2.6.1a)^53^. Transcripts per million (TPM) generation and differential expression analysis was performed on aligned reads to Mus musculus UCSC GRCm38/mm10 reference genome using Illumina RNA-seq differential expression software (DESeq2, v1.0.1)^54^. Significance cut-offs are listed in figure legends for each analysis. GSEA^55,56^ was performed by comparing DKO cells to all other genotypes combined (WT, ACLY KO, ACSS2 KO).

### Immunofluorescence confocal microscopy

Cells were plated on glass coverslips and allowed to adhere for 24 hours. Live cells were then incubated in DMEM + 10% FS with 1 μM MitoTracker Deep Red FM (Thermo) for 30 minutes. Cells were washed with PBS 3 times then fixed with 4% PFA for 30 minutes. After fixation, coverslips were washed 3 times with TBST then blocked with 10% goat serum, 1% BSA, 0.1% gelatin, 22.52 mg/mL glycine, and 0.1% Triton X-100 in TBST for 30 minutes. Coverslips were washed with TBST 3 times then incubated with PDHe1α antibody (Abcam ab110334) for 3 hours. Coverslips were washed with TBST 3 times then incubated with DAPI and donkey anti-rabbit Alexa Flour 488 (Invitrogen #a21206) for 1 hour. Coverslips were washed 3 times then mounted on slides for imaging. Slides were imaged on a Zeiss LSM 880. Z-stacks were compressed into a single plane for representation.

### Statistical Analyses

All analyses were performed using GraphPad Prism or R (RNA sequencing analysis). Statistical test specifics are included in figure legends.

## Author Contributions

Conceptualization: LI, KEW, NWS

Methodology: LI, KEW, NWS

Formal Analysis: LI, ST, CD, TM, NK, AF

Investigation: LI, ST, CD, JD, TM, NK, AF, PN, LR, JS, HA, AC

Visualization: LI

Supervision: KEW, NWS, BC, AA

Writing - original draft: LI, KEW

Writing - review & editing: LI, KEW, NWS, ST, AC, BC, AA, CD, JD, TM, NK, AF, PN, LR, JS, HA

## Data and Material Availability

All data needed to evaluate the conclusions in the paper are present in the paper and/or the Supplementary Materials. RNA sequencing data will be deposited in GEO.

