## Supplemental Figures and Tables for "The carnitine shuttle links mitochondrial metabolism to histone acetylation and lipogenesis"

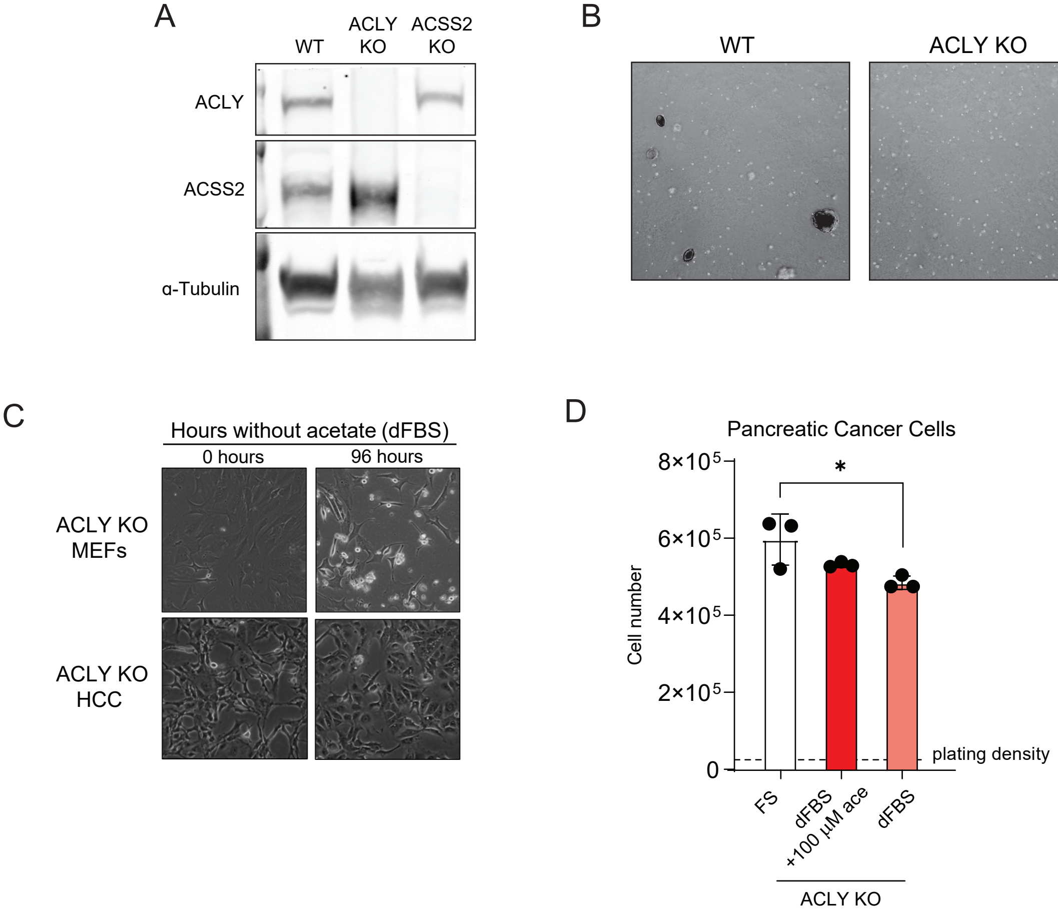


Supplemental Figure 1: ACLY KO cells remain viable without acetate

**Supplemental Figure 1: ACLY KO cells remain viable without acetate**

A) Western blot for ACLY and ACSS2 in WT, ACLY KO, and ACSS2 KO HCC cells.

B) Representative images of soft agar colony formation assay for WT, ACLY KO, and ACSS2 KO HCC cells.

C) Representative images of ACLY KO MEFs (PC9 cells) and ACLY KO HCC cells grown in the absence of acetate for 0 hours and 96 hours. Cells were plated near confluency and monitored for cell death through decreasing confluency over time.

D) Pancreatic cancer ACLY KO cell proliferation in DMEM + 10% FS, DMEM + 10% dFBS, or DMEM + 10% dFBS + 100μM acetate for 72 hours. Statistical significance was calculated by one-way ANOVA

Each point represents a biological replicate and error bars represent standard deviation. *p≤0.05; **p≤0.01; ***p≤0.001; ****p≤0.0001

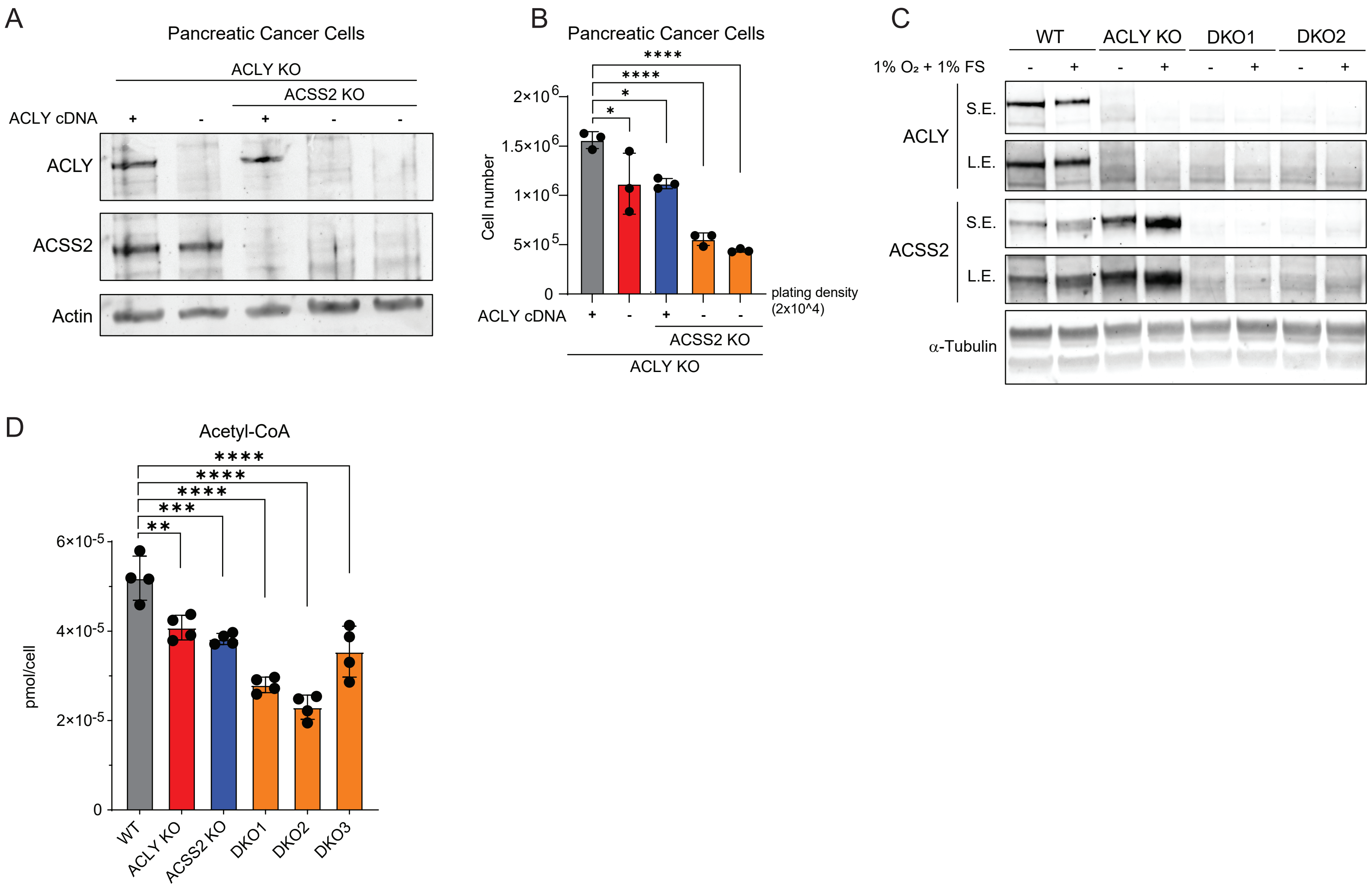


Supplemental Figure 2: DKO cancer cells can proliferate and maintain acetyl-CoA

**Supplemental Figure 2: DKO cancer cells can proliferate and maintain acetyl-CoA**

A) Western blot for ACLY and ACSS2 in Pancreatic cancer ACLY KO cell lines expressing ACLY cDNA or lacking ACSS2 after CRISPR/Cas9 mediated knockout.

B) Proliferation of pancreatic cancer cell lines over 5 days in DMEM + 10% FS. Statistical significance was calculated by one-way ANOVA.

C) Western blot for ACLY and ACSS2 in HCC cells cultured in 10% FS at atmospheric oxygen (-) or 1% FS in 1% oxygen (+) for 24 hours. S.E. short exposure; L.E. long exposure.

D) Whole cell acetyl-CoA measurements in HCC cells cultured in DMEM + 10% FS. Statistical significance was calculated by one-way ANOVA.

Each point represents a biological replicate and error bars represent standard deviation. $^*p \leq 0.05$; $^{**}p \leq 0.01$; $^{***}p \leq 0.001$; $^{****}p \leq 0.0001$

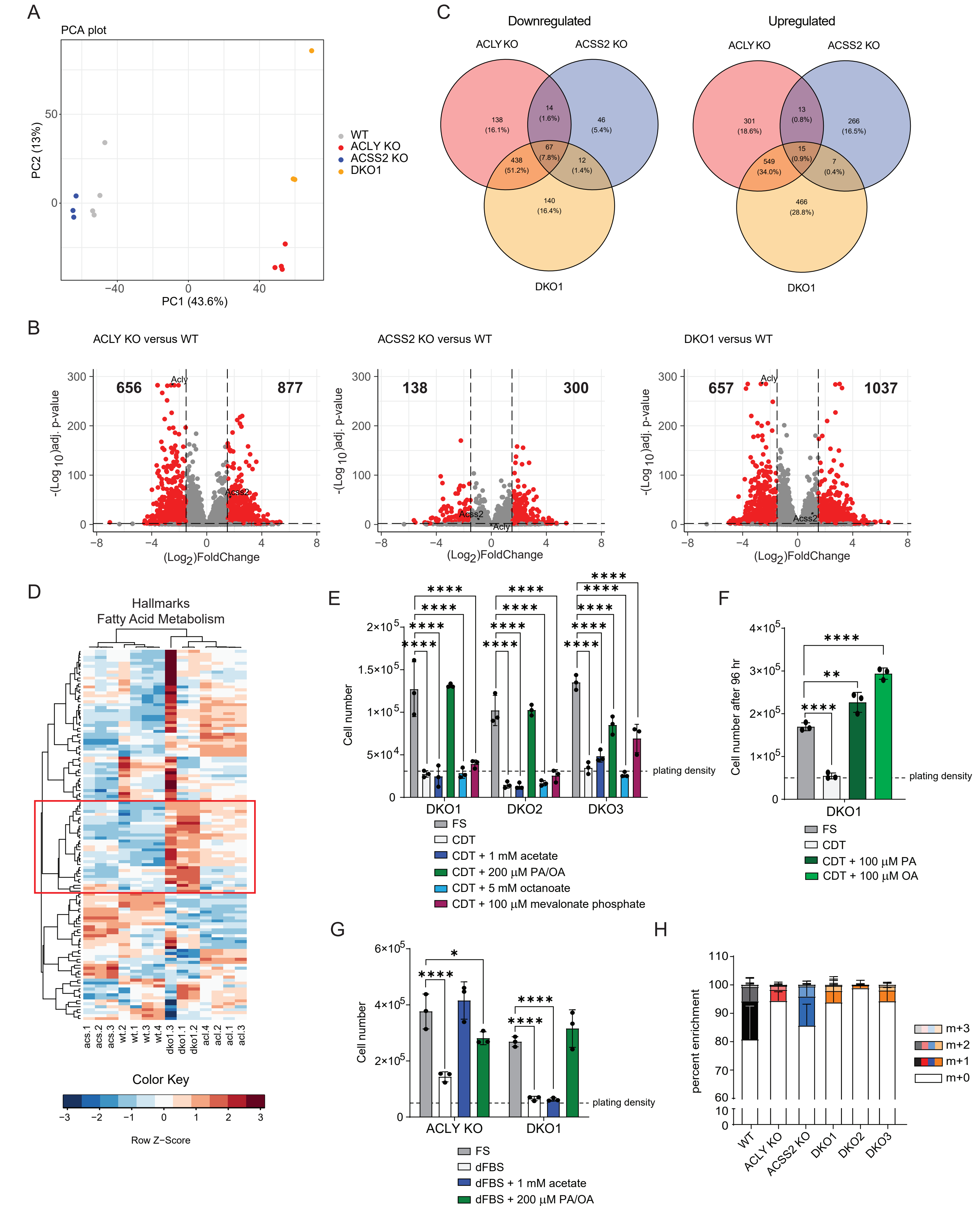


Supplemental Figure 3: Fatty acid metabolism is altered in DKO cells.

**Supplemental Figure 3: Fatty acid metabolism is altered in DKO cells.**

A) Principal component analysis of log2 transformed DESeq counts from RNAseq performed on cells grown in DMEM + 10% FS.

B) Volcano plots showing differentially expressed genes comparing KO genotypes to WT cells. Dots represent individual genes. Red dots are genes with a log2 fold change>1.5 and adjusted p-value<0.01.

C) Venn diagrams comparing the differentially expressed genes (log2 fold change>1.5 and adjusted p-value<0.01) between each KO genotype and WT cells.

D) Heatmap of all genes from the hallmarks fatty acid metabolism gene set. DESeq counts were log2 transformed before clustering. Red box shows the gene cluster expanded in figure 3C.

E) Cell proliferation after 96 hours. Cells were plated in DMEM/F12 media overnight then cultured in DMEM + 10% FS or CDT serum with or without the addition of metabolites. PA/OA is 100 µM of each fatty acid conjugated to BSA (200 µM total). Statistical significance was calculated by two-way ANOVA.

F) Cell proliferation after 96 hours. Cells were plated in DMEM/F12 media for overnight then cultured in DMEM + 10% FS or CDT serum with or without the addition of 100 µM of each fatty acid conjugated to BSA. Statistical significance was calculated by one-way ANOVA.

G) Cell proliferation after 96 hours. Cells were plated in DMEM/F12 media overnight then cultured in DMEM + 10% FS or dFBS serum with or without the addition of metabolites. PA/OA is 100 µM of each fatty acid conjugated to BSA (200 µM total). Statistical significance was calculated by two-way ANOVA.

H) Isotopologue enrichment of palmitate measured by GC-MS. Cells were cultured in DMEM + 10% D2O + 10% FS for 24 hours.

Each point represents a biological replicate and error bars represent standard deviation. *p≤0.05; **p≤0.01; ***p≤0.001; ****p≤0.0001

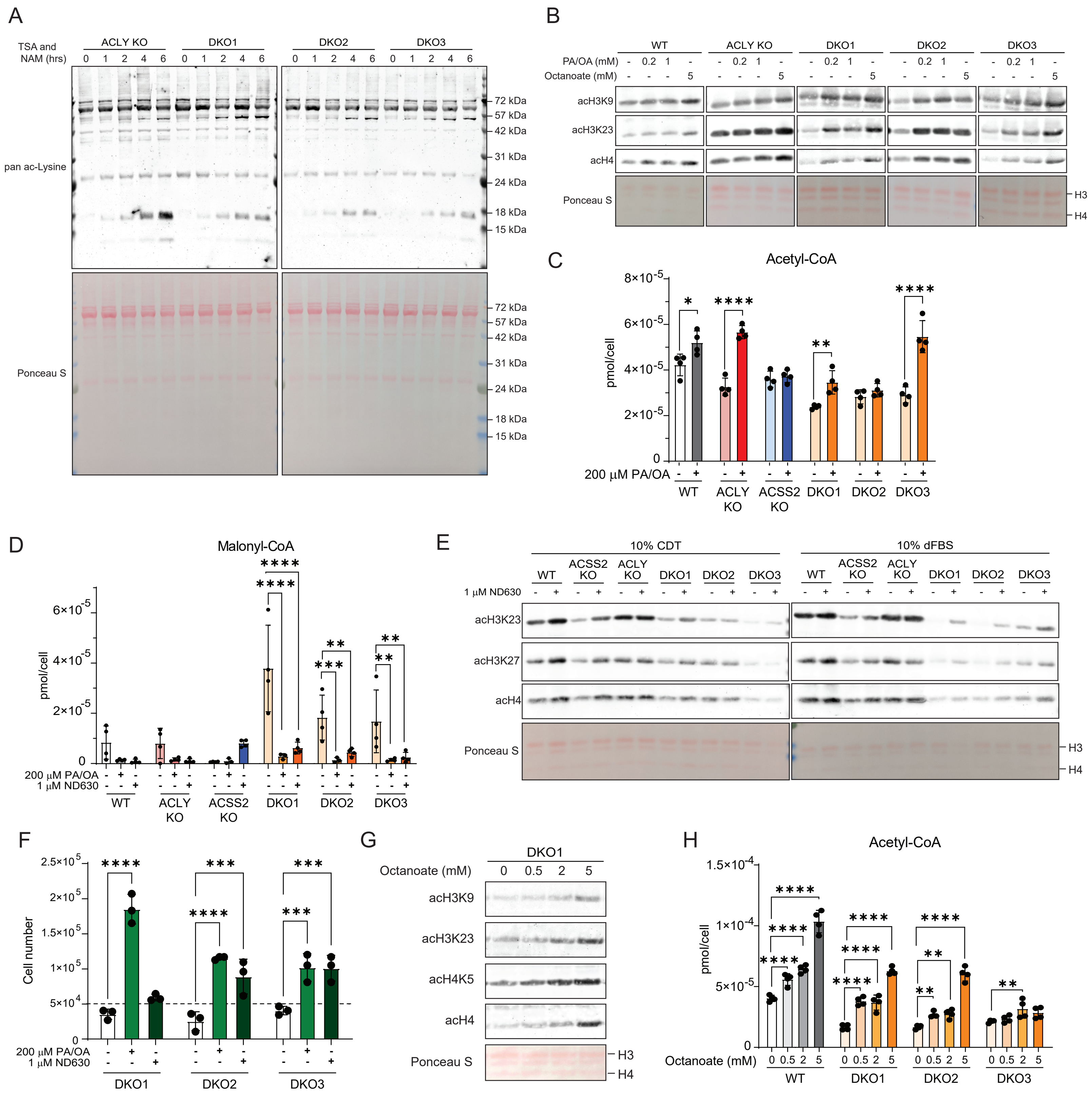


Supplemental Figure 4: Regulation of acetylation by fatty acids may act through acetyl-CoA sparing and ACC inhibition

**Supplemental Figure 4: Regulation of acetylation by fatty acids may act through acetyl-CoA sparing and ACC inhibition**

A) Whole cell protein extract western blot from cells grown in DMEM + 10% FS and treated with 500 nM TSA and 500 μM nicotinamide (NAM) over a time course.

B) Acid extracted histone western blot from cells cultured in DMEM + 10% CDT and supplemented with PA/OA or 5mM octanoate. PA/OA is 100 or 500 μM of each fatty acid conjugated to BSA. Ponceau S stain for total protein in histone extracts used for western blot.

C) Whole cell acetyl-CoA quantitation of cells cultured in DMEM + 10% CDT with or without the addition PA/OA. PA/OA is 100 μM of each fatty acid conjugated to BSA (200 μM total). Statistical significance was calculated by multiple t-tests.

D) Whole cell malonyl-CoA quantitation of cells cultured in DMEM + 10% CDT with or without the addition PA/OA or the ACC inhibitor ND630. PA/OA is 100 μM of each fatty acid conjugated to BSA (200 μM total). Statistical significance was calculated by two-way ANOVA.

E) Acid extracted histone western blot from cells cultured in DMEM + 10% CDT or DMEM + 10% dFBS with or without ND630. Ponceau S stain for total protein in histone extracts used for western blot.

F) Cell proliferation after 96 hours. Cells were plated in DMEM/F12 media for overnight then cultured in DMEM + 10% CDT serum with or without the addition PA/OA or the ACC inhibitor ND630. Statistical significance was calculated by two-way ANOVA.

G) Acid extracted histone western blot from cells cultured in DMEM + 10% FS and octanoate for 24 hours.

H) Whole cell acetyl-CoA quantitation of WT and DKO cells cultured in DMEM + 10% FS supplemented with octanoate for 24 hours. Statistical significance was calculated by two-way ANOVA.

Each point represents a biological replicate and error bars represent standard deviation. *$p \leq 0.05$; **$p \leq 0.01$; ***$p \leq 0.001$; ****$p \leq 0.0001$

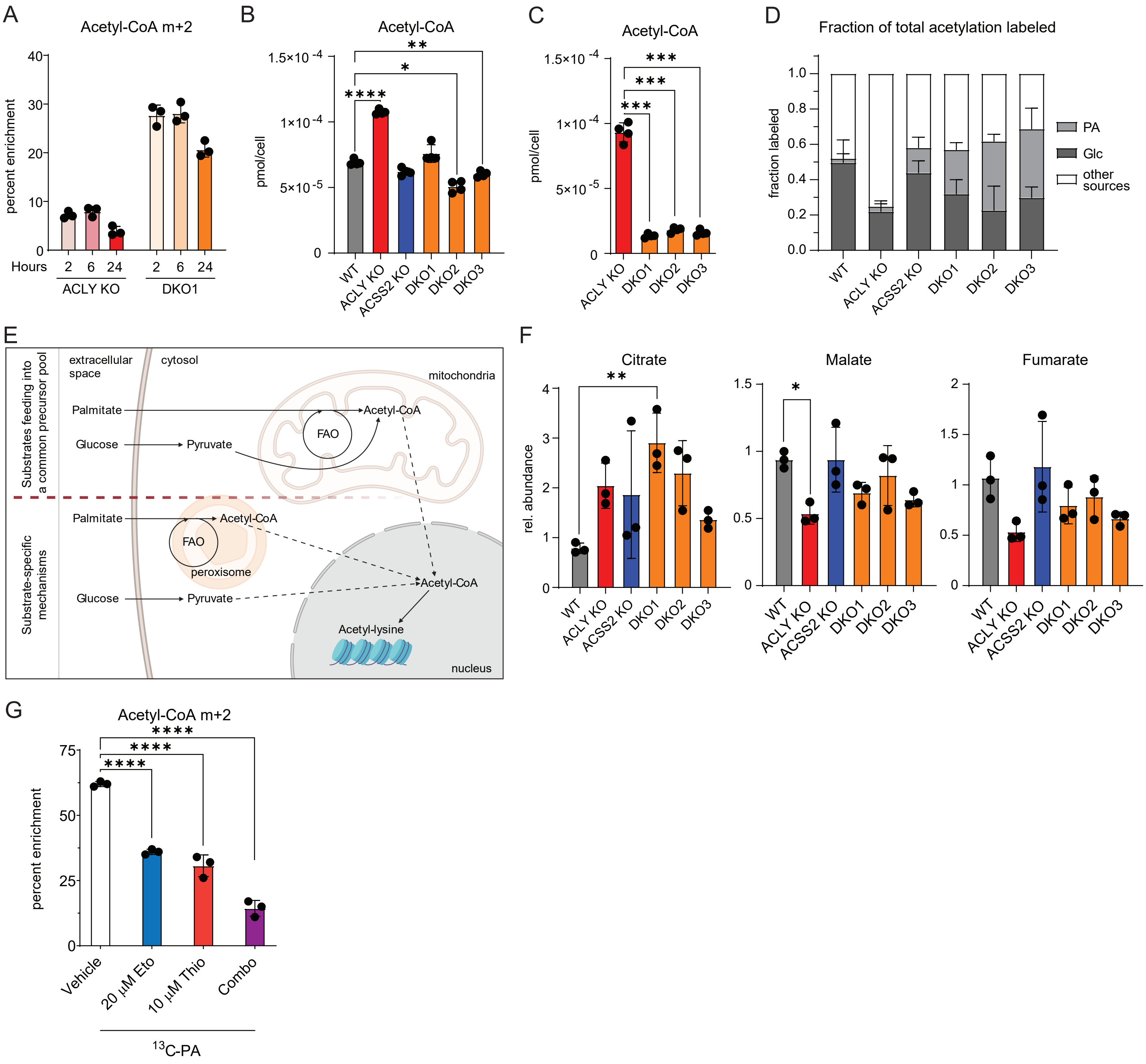


Supplemental Figure 5: Glucose and fatty acid contribution to histone acetylation may be through parallel or convergent pathways

**Supplemental Figure 5: Glucose and fatty acid contribution to histone acetylation may be through parallel or convergent pathways**

A) $^{13}C_{16}$-palmitate tracing into acetyl-CoA analyzed by LC-MS. Cells were cultured in glucose and glutamine free DMEM + 10% CDT supplemented with 4 mM glutamine, 10 mM glucose and 100 μM $^{13}C_{16}$-palmitate conjugated to BSA for 2, 6, or 24 hours.

B) Whole cell acetyl-CoA quantitation in cells grown in glucose and glutamine free DMEM + 10% CDT supplemented with 4 mM glutamine and either 10 mM $^{13}C_6$-glucose and 100 μM palmitate conjugated to BSA or 10 mM glucose and 100 μM $^{13}C_{16}$-palmitate conjugated to BSA for 2 hours. Statistical significance was calculated by one-way ANOVA.

C) Whole cell acetyl-CoA quantitation in cells grown in glucose and glutamine free DMEM + 10% dFBS supplemented with 4 mM glutamine and 10 mM glucose and 100 μM acetate for 6 hours. Statistical significance was calculated by one-way ANOVA.

D) Total of all acetylation measured on acid extracted histone coming from $^{13}C_6$-glucose (Glc) or $^{13}C_{16}$-palmitate (PA) after 24 hours or incubation. Unlabeled acetylation marks are shown as "other sources".

E) Schematic showing potential explanations for glucose and palmitate labeling into acetyl-groups in the nuclear compartment. Top half represents a route for both palmitate and glucose to feed into a precursor acetyl-CoA pool in the mitochondria, bottom half represents two potential separate pathways involving peroxisomal beta-oxidation of fatty acids and pyruvate conversion to acetyl-CoA outside of the mitochondria. Created with BioRender.com.

F) TCA cycle intermediate relative quantitation. Statistical significance was calculated by one-way ANOVA.

G) $^{13}C_{16}$-palmitate tracing into acetyl-CoA analyzed by LC-MS. Cells were cultured in glucose and glutamine free DMEM + 10% CDT supplemented with 4 mM glutamine, 10 mM glucose and 100 μM $^{13}C_{16}$-palmitate conjugated to BSA for 2 hours. Cells were treated for 15 minutes prior to labeling with the CPT1 inhibitor etomoxir (20 μM Eto), the peroxisomal beta-oxidation inhibitor thioridazine (10 μM Thio) or a combination of both drugs (Combo). Statistical significance was calculated by one-way ANOVA.

Each point represents a biological replicate and error bars represent standard deviation. $^{*}p \le 0.05$; $^{**}p \le 0.01$; $^{***}p \le 0.001$; $^{****}p \le 0.0001$

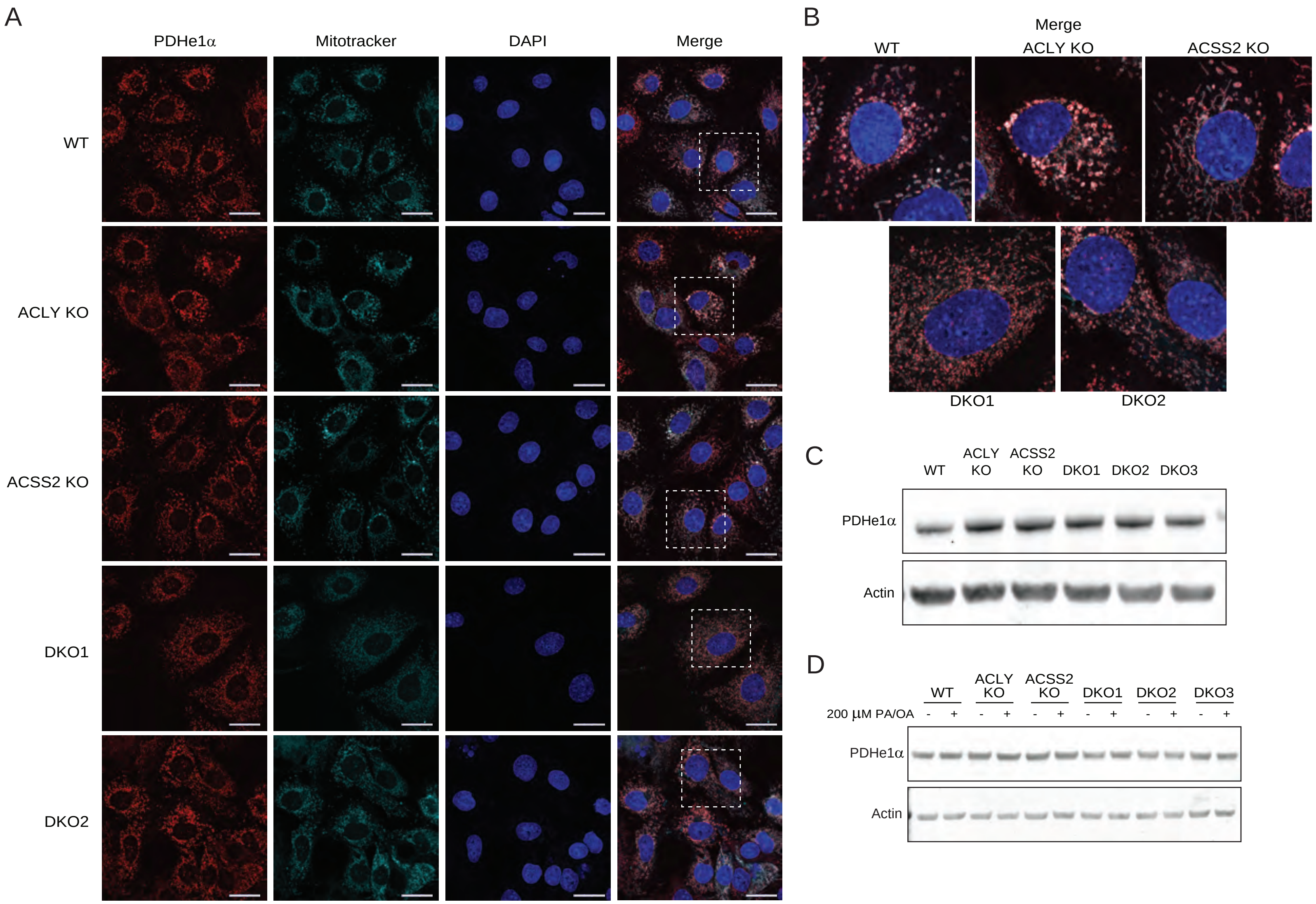


Supplemental Figure 6: PDH co-localizes with the mitochondria and is not regulated at the protein level by ACLY, ACSS2, or fatty acids

**Supplemental Figure 6: PDH co-localizes with the mitochondria and is not regulated at the protein level by ACLY, ACSS2, or fatty acids**

A) Confocal microscopy images of HCC cells cultured in DMEM + 10% FS for 24 hours. Scale bar is 25 µM.

B) Enlarged images from dotted inserts in panel A.

B) Western blot for PDHe1α in whole cell lysates from HCC cells cultured in DMEM + 10% FS for 24 hours.

C) Western blot for PDHe1α in whole cell lysates from HCC cells cultured in DMEM + 10% CDT with or without the addition PA/OA for 24 hours. PA/OA is 100 µM of each fatty acid conjugated to BSA (200 µM total).

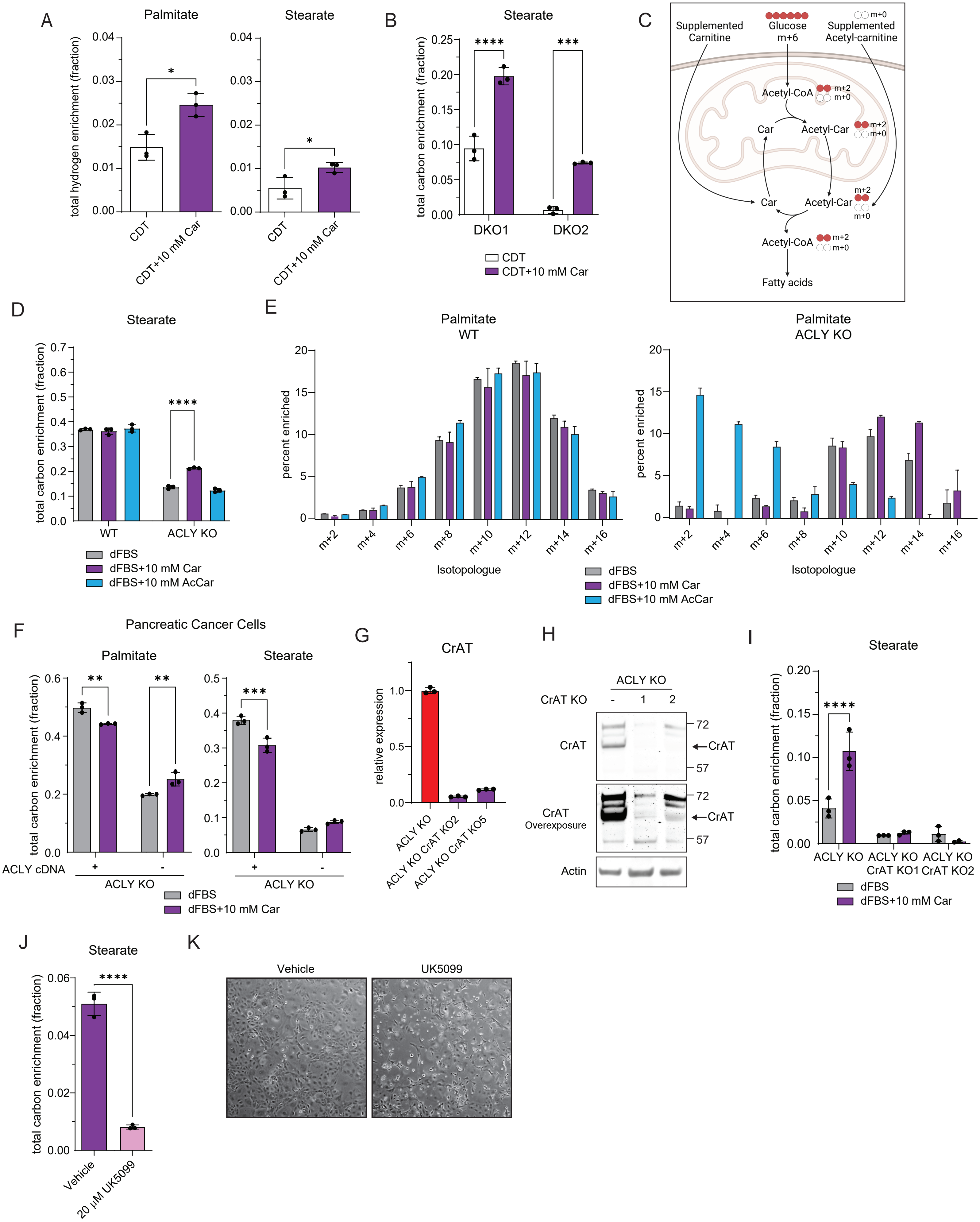


Supplemental Figure 7: Carnitine shuttling increases glucose derived de novo lipogenesis

**Supplemental Figure 7: Carnitine shuttling increases glucose derived de novo lipogenesis**

A) Deuterium tracing into palmitate and stearate measured by GC-MS. Cells were cultured in DMEM + 10% $D_2O$ + 10% CDT, +/- 10 mM carnitine for 24 hours. Statistical significance was calculated by unpaired t-tests.

B) $^{13}C_6$-glucose tracing into stearate measured by GC-MS. Cells were cultured in glucose and glutamine free DMEM + 10% CDT supplemented with 4 mM glutamine and 10 mM $^{13}C_6$-glucose, +/- 10 mM carnitine for 24 hours. Statistical significance was calculated by unpaired t-tests.

C) Schematic showing glucose labeling into fatty acids through the proposed carnitine shuttle and the impact of addition of exogenous carnitine or acetylcarnitine to cells. Created with BioRender.com.

D) $^{13}C_6$-glucose tracing into stearate measured by GC-MS after esterification of fatty acids. Cells were cultured in glucose and glutamine free DMEM + 10% dFBS supplemented with 4 mM glutamine and 10 mM $^{13}C_6$-glucose with or without 10 mM carnitine or 10 mM acetylcarnitine for 24 hours.

E) Full isotopologue distribution of $^{13}C_6$-glucose tracing into palmitate measured by GC-MS. Cells were cultured in glucose and glutamine free DMEM + 10% dFBS supplemented with 4 mM glutamine and 10 mM $^{13}C_6$-glucose with or without 10 mM carnitine or 10 mM acetylcarnitine for 24 hours.

F) $^{13}C_6$-glucose tracing into palmitate and stearate measured by GC-MS in pancreatic cancer cells. Cells were cultured in glucose and glutamine free DMEM + 10% dFBS supplemented with 4 mM glutamine and 10 mM $^{13}C_6$-glucose with or without 10 mM carnitine for 24 hours. Statistical significance was calculated by unpaired t-tests.

G) RT-qPCR analysis of CrAT KO clones. ΔΔCt analysis was performed using actin as the internal control gene and ACLY KO cells as the control condition.

H) Western blot for CrAT in whole cell lysates from HCC cells cultured in DMEM + 10% FS for 24 hours.

I) $^{13}C_6$-glucose tracing into stearate measured by GC-MS after esterification of fatty acids. Cells were cultured in glucose and glutamine free DMEM + 10% dFBS supplemented with 4 mM glutamine and 10 mM $^{13}C_6$-glucose, +/- 10 mM carnitine for 24 hours. Statistical significance was calculated by unpaired t-tests.

J) $^{13}C_6$-glucose tracing into stearate measured by GC-MS after esterification of fatty acids. Cells were cultured in glucose and glutamine free DMEM + 10% dFBS supplemented with 4 mM glutamine and 10 mM $^{13}C_6$-glucose with 10 mM carnitine with vehicle control or 20 μM UK5099 for 48 hours. Statistical significance was calculated by unpaired t-tests.

K) Representative images of ACLY KO cells grown in glucose and glutamine free DMEM + 10% dFBS supplemented with 4 mM glutamine and 10 mM $^{13}C_6$-glucose with 10 mM carnitine with vehicle control or 20 μM UK5099 for 48 hours.

Each point represents a biological replicate and error bars represent standard deviation. *p≤0.05; **p≤0.01; ***p≤0.001; ****p≤0.0001

**Supplemental Table 1: Differentially regulated genes from cluster analysis in Figure 3A**

**Table S1, Related to Figure 3A**

Gene list and cluster ID for WT, ACLY KO, ACSS2 KO, and DKO cells represented on heat map

| Symbol | Cluster |
|---|---|
| Slc6a12 | red |
| Cttnbp2 | red |
| Elfn2 | red |
| Csmd1 | red |
| Taf7l | red |
| Crhbp | red |
| Ndst4 | red |
| Trpm5 | red |
| Ly6i | red |
| Toporsl | red |
| Npc1l1 | red |
| Ak8 | red |
| Slc47a1 | red |
| Myoz2 | red |
| Pdcd1lg2 | red |
| Megf11 | red |
| Saa3 | red |
| Naip6 | red |
| Sprr2i | red |
| D830026I12Rik | red |
| 1700019E08Rik | red |
| Chgb | red |
| A630095N17Rik | red |
| Hs3st3a1 | red |
| Adgrl4 | red |
| Pla2r1 | red |
| Dthd1 | red |
| Arsj | red |
| Acaa1b | red |
| 4930592C13Rik | red |
| Gm1965 | red |
| Gm10451 | red |
| 5430428K19Rik | red |
| Otogl | red |
| 5430425K12Rik | red |
| Apol6 | red |
| Gm973 | red |
| A230001M10Rik | red |
| Gm13031 | red |
| Arhgef6 | red |
| Casq2 | red |
| C330024C12Rik | red |
| Plek | red |
| Fam83a | red |

| | |
|---|---|
| Gm1110 | red |
| 4930503B20Rik | red |
| Sox5os3 | red |
| 5430427O19Rik | red |
| Fcgr2b | red |
| Ceacam10 | red |
| Tmem132e | red |
| Prl3d2 | red |
| Gm1322 | red |
| Eef1a2 | red |
| Cd177 | red |
| Tigd4 | red |
| Itgam | red |
| Trpm2 | red |
| Gm8267 | red |
| Gm4861 | red |
| 4932414N04Rik | red |
| 4933405E24Rik | red |
| D930007P13Rik | red |
| Alpk2 | red |
| Gm28453 | red |
| 2810405F15Rik | red |
| Fgl1 | red |
| Cryba4 | red |
| BC061237 | red |
| 9230020A06Rik | red |
| Pzp | red |
| Gm4371 | red |
| Alpi | red |
| Serpinb9f | red |
| 4930509E16Rik | red |
| Lypd6 | red |
| U90926 | red |
| Kcnn4 | red |
| Sult1d1 | red |
| Atp2c2 | red |
| Gsg1l | red |
| Coro1a | red |
| Nps | red |
| 1700018A04Rik | red |
| Ccdc155 | red |
| Brinp3 | red |
| Cd28 | red |
| 5330434G04Rik | red |
| Hgf | red |
| Gm2721 | red |
| Nhlrc4 | red |
| Acot12 | red |

| | |
|---|---|
| Cngb1 | red |
| Catip | red |
| Gbx1 | red |
| Olfr875 | red |
| Mill2 | red |
| Olfr225 | red |
| Tnnt3 | red |
| Phf11b | red |
| Mmp20 | red |
| Gm11529 | red |
| Slc35f1 | red |
| Aim2 | red |
| 1700092C10Rik | red |
| D030045P18Rik | red |
| Cntn3 | red |
| Umod | orange |
| Hist1h1t | orange |
| 1600019K03Rik | orange |
| Kctd14 | orange |
| Hist1h2ad | orange |
| Cts3 | orange |
| Apof | orange |
| Mtus2 | orange |
| Stk33 | orange |
| Sprn | orange |
| Lbx1 | orange |
| Tro | orange |
| 1700113B09Rik | orange |
| Gm5535 | orange |
| Ugt1a7c | orange |
| 1700012B07Rik | orange |
| Prelid2 | orange |
| LOC108168459 | orange |
| Nap1l3 | orange |
| C130026I21Rik | orange |
| Sema5b | orange |
| 44631 | orange |
| Ttc29 | orange |
| Hspb7 | orange |
| Gm12238 | orange |
| Bank1 | orange |
| Lyzl1 | orange |
| Cmtm5 | orange |
| Uba1y | orange |
| Wnt1 | orange |
| Csf2rb2 | orange |
| 9330178D15Rik | orange |
| Serpina3n | orange |

| | |
|---|---|
| 4921511I17Rik | orange |
| Unc5a | blue |
| Elavl2 | blue |
| Frzb | blue |
| Dlx6 | blue |
| Sim1 | blue |
| Shank3 | blue |
| Wipf3 | blue |
| Col23a1 | blue |
| Adamts13 | blue |
| 1700007K13Rik | blue |
| Dlx6os1 | blue |
| B4galnt2 | blue |
| H2-Q1 | blue |
| BB218582 | blue |
| Asxl3 | blue |
| Tubb2a-ps2 | light grey |
| Eif4e3 | light grey |
| Hmga2-ps1 | light grey |
| Zfp354b | light grey |
| Itgb8 | light grey |
| Gm15050 | light grey |
| Dusp23 | light grey |
| Otulinl | light grey |
| Cbr3 | light grey |
| Sfrp2 | light grey |
| Pou4f1 | light grey |
| Hddc3 | light grey |
| Ctsf | light grey |
| Htr1b | light grey |
| H60b | light grey |
| Phlda2 | light grey |
| Ccdc30 | light grey |
| Tbx2 | light grey |
| Ms4a4b | light grey |
| Cdh15 | light grey |
| Crmp1 | light grey |
| Triqk | light grey |
| Tmem121 | light grey |
| Serpina3j | light grey |
| 1700034H15Rik | light grey |
| Cfap100 | light grey |
| 1700012C14Rik | light grey |
| Zfp783 | light grey |
| Nfasc | light grey |
| Rasgef1a | light grey |
| Tlr6 | light grey |
| Mir3063 | light grey |

| | |
|---|---|
| Hmx3 | light grey |
| Gm13648 | light grey |
| Ptgdr2 | light grey |
| Cln3 | light grey |
| Fads6 | dark grey |
| Amph | dark grey |
| Lhfpl4 | dark grey |
| A030001D20Rik | dark grey |
| Tdh | dark grey |
| Ror2 | dark grey |
| Ppp1r14a | dark grey |
| Gstm4 | dark grey |
| Sp5 | dark grey |
| Galm | dark grey |
| Hexa | dark grey |
| Gm8773 | dark grey |
| Dusp9 | dark grey |
| Rhoj | dark grey |
| Misp | dark grey |
| Sowahb | dark grey |
| Cdh26 | dark grey |
| Mansc1 | dark grey |
| Rgs4 | dark grey |
| Drd3 | dark grey |
